# Controlled Induction of ALK1 in *ACVRL1*-null hiPSC-derived Endothelial Cells Provides Insight into Organ-Restricted AVMs in HHT

**DOI:** 10.1101/2025.08.22.671484

**Authors:** Sebastiaan J. van Kampen, Albert Blanch-Asensio, James L. Gallant, Babet van der Vaart, Christian M.A.H. Freund, Richard P. Davis, Christine L. Mummery, Valeria V. Orlova

## Abstract

Genetic alterations in activin receptor-like kinase 1 (*ACVRL1*, ALK1) are linked to hereditary hemorrhagic telangiectasia (HHT), a condition characterized by hemorrhages, arteriovenous malformations (AVMs) and endothelial cell (EC) dysfunction. Haploinsufficiency is considered the central disease-driving mechanism, but it remains unclear why vascular lesions are tissue-restricted. To investigate whether *ACVRL1* gene dosage in human ECs plays a role in organ-specific susceptibility, we developed a tunable human *in vitro* model by deleting *ACVRL1* in human induced pluripotent stem cells (hiPSCs), and then reintroduced wild-type *ACVRL1* under control of a doxycycline-inducible promoter. This enabled temporal, dose-dependent induction of ALK1 expression in ECs, allowing definition of threshold levels required to restore endothelial function and support vascular homeostasis. ALK1-deficient hiPSC-ECs displayed disrupted SMAD1/5 signaling, hyperproliferation, altered tip/stalk cell specification, and dysregulated transcriptional programs. Even low-level reinduction of ALK1 was sufficient to restore endothelial function, indicating threshold-dependence for ALK1 in vascular homeostasis. These data indicated that one mechanism underlying tissue-restricted prevalence of AVMs in vascular beds with low basal *ACVRL1* expression, such as the liver, is a minimal requirement for ALK1 in sustaining EC functionality. This tunable human platform offers a powerful tool for dissecting HHT pathobiology and a platform for identifying strategies to restore ALK1 signaling therapeutically in affected tissues.

## INTRODUCTION

Hereditary hemorrhagic telangiectasia (HHT) is an inherited vascular disorder characterized by telangiectasias and arteriovenous malformations (AVMs), often affecting the skin, lungs, liver, brain, and gastrointestinal tract (Komiyama et al., 2014; Kuehl et al., 2005; McDonald et al., 2000; Sabba et al., 2007; Sanchez-Martinez et al., 2020). Telangiectasias frequently occur on the lips, tongue, and ocular mucosa, often leading to recurrent epistaxis and iron deficiency anemia due to chronic blood loss (McAllister et al., 1994; Shovlin et al., 2000). AVMs on the other hand can cause serious complications through abnormal arteriovenous shunting in internal organs. Current treatment options focus on reducing symptom burden rather than curing the disease due to a paucity of therapeutic targets (Al-Samkari, 2024), although several new drugs are currently in clinical trial (Al Tabosh et al., 2024).

The majority of HHT patients carry heterozygous mutations in either *ACVRL1* or *ENG*, which encode transforming growth factor beta (TGF-β) / bone morphogenetic protein (BMP) receptors activin receptor-like kinase 1 (ALK1) and endoglin (ENG), respectively, both essential for vascular homeostasis (McDonald et al., 2020). Less commonly, mutations in *SMAD4*, a downstream transcriptional mediator in this pathway, result in SMAD4-related HHT, often accompanied by juvenile polyposis. In ALK1-and ENG-related HHT, haploinsufficiency is widely regarded as the primary pathogenic cause, disrupting essential angiogenic pathways such as SMAD-mediated transcription and VEGF signaling in endothelial cells (ECs). These disruptions ultimately promote abnormal endothelial proliferation, migration, defective vessel maturation, and increased vascular fragility (Ahmed et al., 2023; David et al., 2007; Kerr et al., 2015; Larrivee et al., 2012; Scharpfenecker et al., 2007; Schmid et al., 2023; Shao et al., 2009; Thalgott et al., 2018; Upton et al., 2009).

In ALK1-related HHT, patients inherit a heterozygous germline loss-of-function mutation in *ACVRL1*, yet AVMs manifest in a tissue-restricted manner, with the liver being one of the most commonly affected organs (Komiyama et al., 2014; Kuehl et al., 2005; McDonald et al., 2000; Sabba et al., 2007; Sanchez-Martinez et al., 2020). This focal manifestation cannot be fully explained by haploinsufficiency alone. Instead, a two-hit model has been proposed, in which somatic inactivation of the remaining functional allele leads to biallelic loss in localized EC populations (Choi et al., 2012; Shaligram et al., 2020). Recent studies of patient-derived AVMs have provided direct evidence of such somatic second-hit events, reinforcing this model (DeBose-Scarlett et al., 2024; Snellings et al., 2019). In addition to genetic inactivation, emerging hypotheses suggest that tissue-specific differences in basal *ACVRL1* and *ENG* expression may further contribute to regional vulnerability. While comparative *ACVRL1* expression data are currently limited to murine EC single-cell transcriptomic datasets (Kalucka et al., 2020), expression-dependent vulnerability has been more clearly demonstrated in ENG-related HHT in mice (Galaris et al., 2021). Together these observations support a model in which both somatic inactivation and tissue-specific variation in expression contribute to the organ-selective susceptibility to AVM formation in HHT.

Nevertheless, the pathomolecular mechanisms driving tissue-specific vulnerability in HHT remain incompletely understood, partly due to the lack of robust preclinical models that faithfully replicate disease pathology. To address this, we developed a tunable human *in vitro* platform to investigate ALK1 dosage effects. We used CRISPR-Cas9 to generate *ACVRL1*-null human induced pluripotent stem cells (hiPSCs) and reintroduced wild-type *ACVRL1* under control of the TRE3G-doxycycline-inducible promoter. This system allows temporal, dose-dependent induction of ALK1 in hiPSC-derived endothelial cells (hiPSC-ECs). Partial expression of ALK1 restored endothelial function in a dose-dependent manner, with recovery of SMAD1/5 signaling defects, hyperproliferation, tip/stalk cell specification, and transcriptional dysregulation. This suggested a threshold-dependent requirement for ALK1 in maintaining EC integrity. These data support a threshold-based model of ALK1 function in ECs and offers a mechanistic framework for understanding how partial loss of signaling leads to tissue-selective vascular malformations in HHT.

## RESULTS

### Generation of a human *in vitro* platform to model ALK1 dosage effects

HHT patient-derived hiPSC lines typically carry a single heterozygous loss-of-function mutation with variable loss of protein, thereby limiting their use in modelling tissue-specific differences in basal *ACVRL1* expression levels (Kim et al., 2024; Orlova et al., 2022). To address this, we aimed to establish a complete *ACVRL1* knockout in a hiPSC line carrying a STRAIGHT-IN v2 landing pad (LP) at the *AAVS1* locus. The STRAIGHT-IN platform facilitates efficient and site-specific integration of large DNA payloads (Blanch-Asensio et al., 2022; Blanch-Asensio et al., 2023). To create the *ACVRL1*-null line, STRAIGHT-IN v2 hiPSCs were transfected simultaneously with Cas9-ribonucleoprotein complexes containing one of three single guide RNAs targeting exon 3 of *ACVRL1*. Edited clones were identified by PCR amplification followed by Sanger sequencing (Figure 1A) and confirmed by droplet digital PCR (ddPCR; Figure S1A). Next, to enable controlled *ACVRL1* expression, we integrated a doxycycline-inducible, all-in-one Tet-On 3G *ACVRL1* expression cassette which also contained a 3xNLS-Tdtomato reporter into the STRAIGHT-IN LP in *ACVRL1*-null hiPSCs (Figure 1B). The same cassette was also integrated into *ACVRL1* wild-type hiPSCs to serve as a genetically-matched control. Following transfection, cells were subjected to zeocin selection to enrich for cells that correctly integrated the DNA payload. Genomic DNA from both selected and non-selected populations was analyzed by ddPCR to detect *attP* and *attR* recombination sites (Figure S1B). The presence of *attR* in most zeocin-selected hiPSCs confirmed successful monoallelic integration of the DNA donor vector. To remove auxiliary sequences flanking the integrated cassette, we transfected the cells with a Cre recombinase expression plasmid, followed by puromycin selection (Figure 1B). Copy number analysis using *RPP30* as the reference gene and *BleoR*, within the auxiliary sequences as the target gene, indicated efficient excision of the auxiliary elements in nearly all cells (Figures S1C and S1D). Both the wild-type and *ACVRL1*-null hiPSC lines harboring the doxycycline-inducible *ACVRL1* cassette showed expression of markers of the undifferentiated state NANOG, OCT3/4, and SSEA-4 (Figure S1E). Here after, we refer to the genetically modified lines as Control (*ACVRL1* wild-type + doxycycline-inducible *ACVRL1*) and ALK1 KO/KO (*ACVRL1*-null + doxycycline-inducible *ACVRL1*).

**Figure 1.**
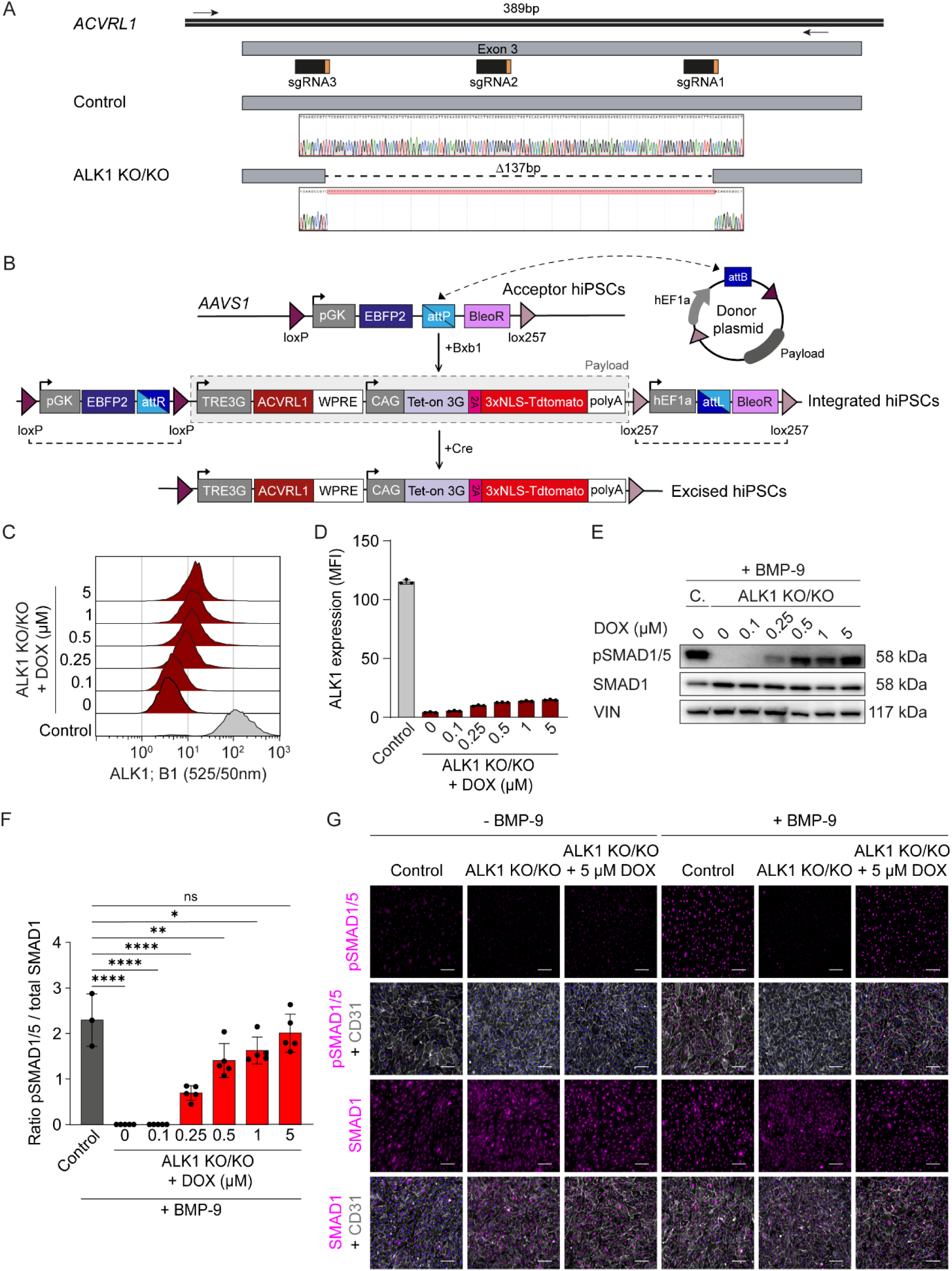
Doxycycline-induced ALK1 expression restores SMAD1/5 activation defects in ALK1 knockout hiPSC-ECs in a dose-dependent manner. (**A**) Schematic depicting exon three of the *ACVRL1* genomic locus which was genetically modified using three single guide RNAs (sgRNA). Selected clones were modified by sgRNAs 1 and 3. Representative Sanger sequencing traces of control and ALK1 knockout hiPSCs. Black arrows indicate the primers used to amplify the targeted region (389bp). (**B**) Schematic of the integration of the doxycycline-inducible *ACVRL1* donor plasmid into the *attP* site present in the *AAVS1* locus of ALK1 knockout hiPSCs and subsequent Cre-mediated excision of the auxiliary elements. (**C**) Representative flow cytometry analysis of ALK1 in control and ALK1 knockout hiPSC-ECs either treated with DMSO or varying concentrations of doxycycline for 72 hours. (**D**) Quantification of surface expression levels of ALK1. N= 3 (independent batches of hiPSC-ECs). (**E-F**) Western blot analysis of extracts from serum starved control and ALK1 knockout hiPSC-ECs stimulated with 10 pg/mL BMP-9 for 45min. ALK1 knockout ECs were treated for a total of 72 hours with the indicated doxycycline concentrations. (**E**) Representative Western blots for pSMAD1/5, total SMAD1, and Vinculin (VIN). (**F**) Quantification of pSMAD1/5 over total SMAD1 ratios. N= 3 (independent batches of hiPSC-ECs); n= 1-3 (one to three technical replicates per batch). (**G**) Representative immunofluorescence images showing either pSMAD1/5 or total SMAD1 levels (magenta), CD31 (gray), and Hoechst (blue) in serum starved control and ALK1 knockout hiPSC-ECs stimulated with DMSO or 10 pg/mL BMP-9 for 45min. Knockout ECs were additionally treated with 5µM doxycycline for 72 hours. Scale bars= 100 μm. Data plotted as mean ±SD. Ordinary one-way ANOVA with Dunnett’s multiple comparisons test (**F**). * p < 0.05; ** p < 0.01; **** p < 0.0001; ns, not significant. DOX, doxycycline; MFI, mean fluorescence intensity; sgRNA, single guide RNA.

### ALK1 induction in ALK1 KO hiPSC-ECs leads to dose-dependent recovery of SMAD1/5 signaling

To investigate the molecular consequences of *ACVRL1* loss in hiPSC-ECs, we first analyzed surface expression levels of ALK1 protein by flow cytometry (Figure S2A). As expected, ALK1 KO/KO hiPSC-ECs showed no detectable ALK1 expression. We next examined if doxycycline could restore ALK1 expression in a dose-dependent manner. ALK1 KO/KO hiPSC-ECs were treated with increasing concentrations of doxycycline for 72 hours, and ALK1 levels quantified by flow cytometry (Figures 1C and 1D). Doxycycline treatment led to a dose-dependent induction of ALK1 expression. However, even at the highest concentration tested (5 μM), ALK1 levels remained ∼7.8 fold lower compared to control cells (Average MFI: control = 115 vs. ALK1 KO/KO + 5 μM = 14.8). Doxycycline concentrations above 5 μM adversely affected cell growth (data not shown), and therefore were not investigated further. Under physiological conditions, ALK1 binds to BMP-9 with high affinity, resulting in the phosphorylation of SMAD1 and SMAD5 (David et al., 2007). To assess whether the activation of SMAD1/5 is compromised in ALK1 KO/KO hiPSC-ECs, we stimulated serum-starved control and knockout cells with varying concentrations of BMP-9 for 45 minutes. Western blot analysis indicated increasing levels of activated SMAD1/5 in control cells, whereas in the absence of doxycycline ALK1 KO/KO hiPSC-ECs were unable to activate SMAD1/5 upon BMP-9 stimulation (Figures S2B and S2C). Interestingly, regardless of BMP-9 presence, total SMAD1 levels were approximately two-fold higher in ALK1 KO/KO hiPSC-ECs compared to control cells (Figure S2D). To explore the extent to which doxycycline-induced ALK1 in ALK1 KO/KO hiPSC-ECs can rescue SMAD1/5 activation defects, we first treated knockout cells with varying concentrations of doxycycline for 72 hours, followed by BMP-9 (10 pg/mL) stimulation for 45 minutes. Western blot (Figures 1E and 1F) and immunofluorescence (Figure 1G) analyses of pSMAD1/5 showed near-complete restoration when treated with 5 μM of doxycycline (Average pSMAD1/5 over SMAD1 ratio: control = 2.29 vs. ALK1 KO/KO + 5 μM = 2.01), despite the low levels of ALK1 induction (Figure 1D). In summary, the progressive induction of ALK1 in ALK1 KO/KO hiPSC-ECs effectively restores the signaling defects associated with SMAD1/5 activation.

### Loss of ALK1 compromises BMP-9 responsiveness in hiPSC-ECs

To better understand the molecular changes underlying the observed SMAD-signaling defects, we conducted bulk mRNA sequencing on both control and ALK1 KO/KO hiPSC-ECs, either unstimulated or stimulated with 1 ng/mL BMP-9 for 12 hours. BMP-9 stimulation of control ECs resulted in a large transcriptional shift, whereas stimulated knockout hiPSC-ECs showed only minor changes compared to their unstimulated counterpart (Figure 2A). These findings were independent of the experimental batch (Figure 2B) or the clonal origin of the lines (Figure 2C). Subsequently, we identified 379 differentially expressed genes (DEGs; logCPM > 1, log2FC > 1 or <-1, and p-adj < 0.05) comparing stimulated control hiPSC-ECs with their unstimulated state (Figure 2D), whereas only 63 DEGs were observed for BMP-9 stimulated knockout ECs compared to baseline (Figure 2E). When comparing ALK1 KO/KO hiPSC-ECs with control cells, both stimulated with BMP-9, we identified 716 DEGs, of which 376 genes were downregulated and 340 upregulated (Figure 2F). Gene Ontology (GO) analysis of the upregulated genes revealed chromosome segregation (GO: 0007059) and cell division (GO: 0051301) as significantly enriched terms, suggesting enhanced proliferation of ALK1 KO/KO hiPSC-ECs (Figure 2G). The enriched GO terms for the downregulated genes included those associated with extracellular matrix (GO: 0005576, 0009986, 0005615) and cell adhesion (GO: 0007155), indicating a disruption in cell-cell and cell-matrix adhesion that could have an impact on vascular integrity (Figure 2H). Collectively, these findings illustrate that ALK1 KO/KO hiPSC-ECs fail to respond adequately to BMP-9.

**Figure 2.**
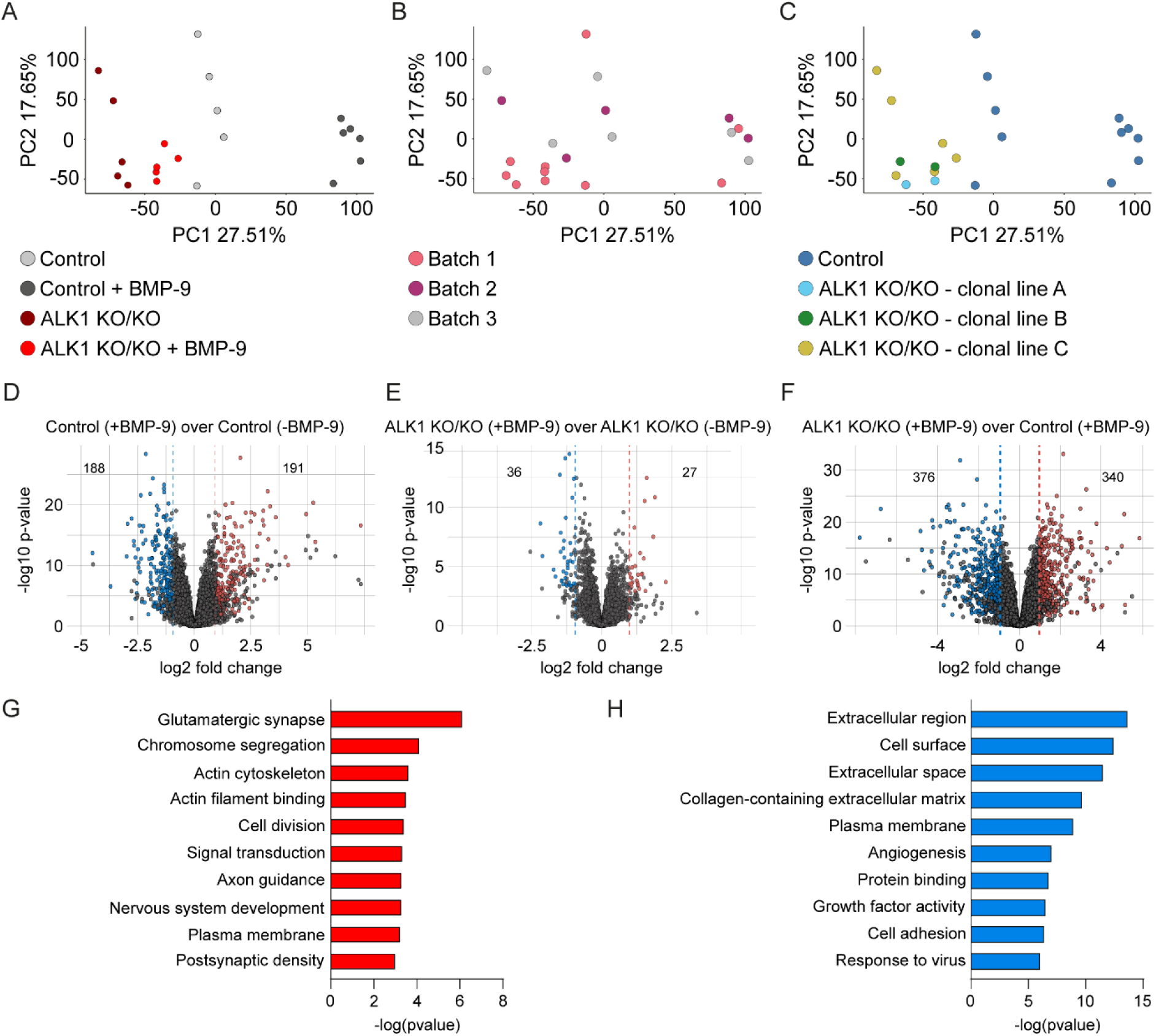
ALK1 knockout hiPSC-ECs are unresponsive to BMP-9 stimulation. (**A-H**) Transcriptomic analysis of control and ALK1 knockout hiPSC-ECs under baseline conditions or after 1 ng/mL BMP-9 stimulation for 12 hours. (**A-C**) Principal-component (PC) plots depicting the experimental conditions (**A**), batch-origin (**B**) and genetic-origin (**C**) of the samples. (**D-F**) Volcano plots showing the differentially up-(red dots) and downregulated (blue dots) genes (logCPM > 1, log2FC > 1 or <-1, and p-adj < 0.05) for the indicated comparisons. (**G-H**) Gene ontology analysis on the up-(**G**) and downregulated (**H**) genes shown in (**F**).

### Gradual ALK1 induction in ALK1 knockout ECs restores gene networks related to SMAD-signaling and cell cycling

To evaluate whether doxycycline-induced ALK1 in ALK1 KO/KO hiPSC-ECs could rescue the disrupted gene networks, we performed bulk mRNA sequencing on control and knockout hiPSC-ECs that were either untreated or exposed to different concentrations of doxycycline for 72 hours and cultured in the absence or presence of 1 ng/mL of BMP-9 for 12 hours. We initially examined whether doxycycline induced *ACVRL1* expression levels in ALK1 KO/KO hiPSC-ECs and found numerous reads mapping to the *ACVRL1* locus (data not shown). Accordingly, we filtered and quantified all reads stemming from the deleted region in exon 3 of *ACVRL1*, which could only originate from the exogenous *ACVRL1* gene inserted in the *AAVS1* locus (Figure 1B). This revealed dose-dependent induction of *ACVRL1* in doxycycline-treated knockout cells, albeit at lower levels than in control cells (Figure 3A), supporting our observation of relatively low levels of surface ALK1 in these cells (Figure 1D). We then explored whether the induced levels of *ACVRL1* led to global transcriptomic changes. Knockout cells treated with doxycycline and stimulated with BMP-9 showed a dose-dependent alteration in their transcriptomic profile, whereas BMP-9 stimulated doxycycline-treated control cells showed only minimal changes (Figure 3B). The experimental batch (Figure 3C) or the clonal origin of the lines (Figure 3D) did not influence the transcriptomic evaluations. Next, we sought to identify all DEGs rescued upon doxycycline-induced ALK1 expression by comparing BMP-9 stimulated knockout hiPSC-ECs with control cells. For this, we compared all knockout subgroups treated with varying concentrations of doxycycline with control cells, reasoning that the DEGs unique to the “BMP-9 stimulated ALK1 knockout hiPSC-ECs vs control” comparison were rescued upon reinduction of ALK1. This approach identified 126 up-(Figure 3E) and 116 downregulated (Figure 3F) DEGs. Semi-supervised clustering of the rescued upregulated DEGs revealed a gene network that, in the absence of BMP-9, was modestly upregulated in ALK1 KO/KO hiPSC-ECs. Furthermore, BMP-9 stimulation did not change the transcriptional pattern of these DEGs in knockout cells, whereas this gene network became suppressed in control cells (Figure 3G). Importantly, doxycycline-induced ALK1 restored this gene network in a dose-dependent manner. Semi-supervised clustering of the rescued downregulated DEGs displayed an inactive gene network in the absence of BMP-9, which became activated in control hiPSC-ECs upon BMP-9 stimulation (Figure 3H). Conversely, BMP-9 stimulated knockout hiPSC-ECs were unable to activate this gene network, although doxycycline-induced ALK1 gradually restored this capacity (Figure 3H). GO analysis of the rescued upregulated DEGs revealed enrichment of terms related to mitosis and cell division (Figure 3I), whereas the rescued downregulated DEGs were associated with TGF-β/BMP biology (Figure 3J). Together, expression of ALK1 in ALK1 KO/KO hiPSC-ECs gradually restored transcriptional responses to BMP-9 stimulation, thereby rescuing gene networks linked to TGF-β/BMP biology and cell cycle progression.

**Figure 3.**
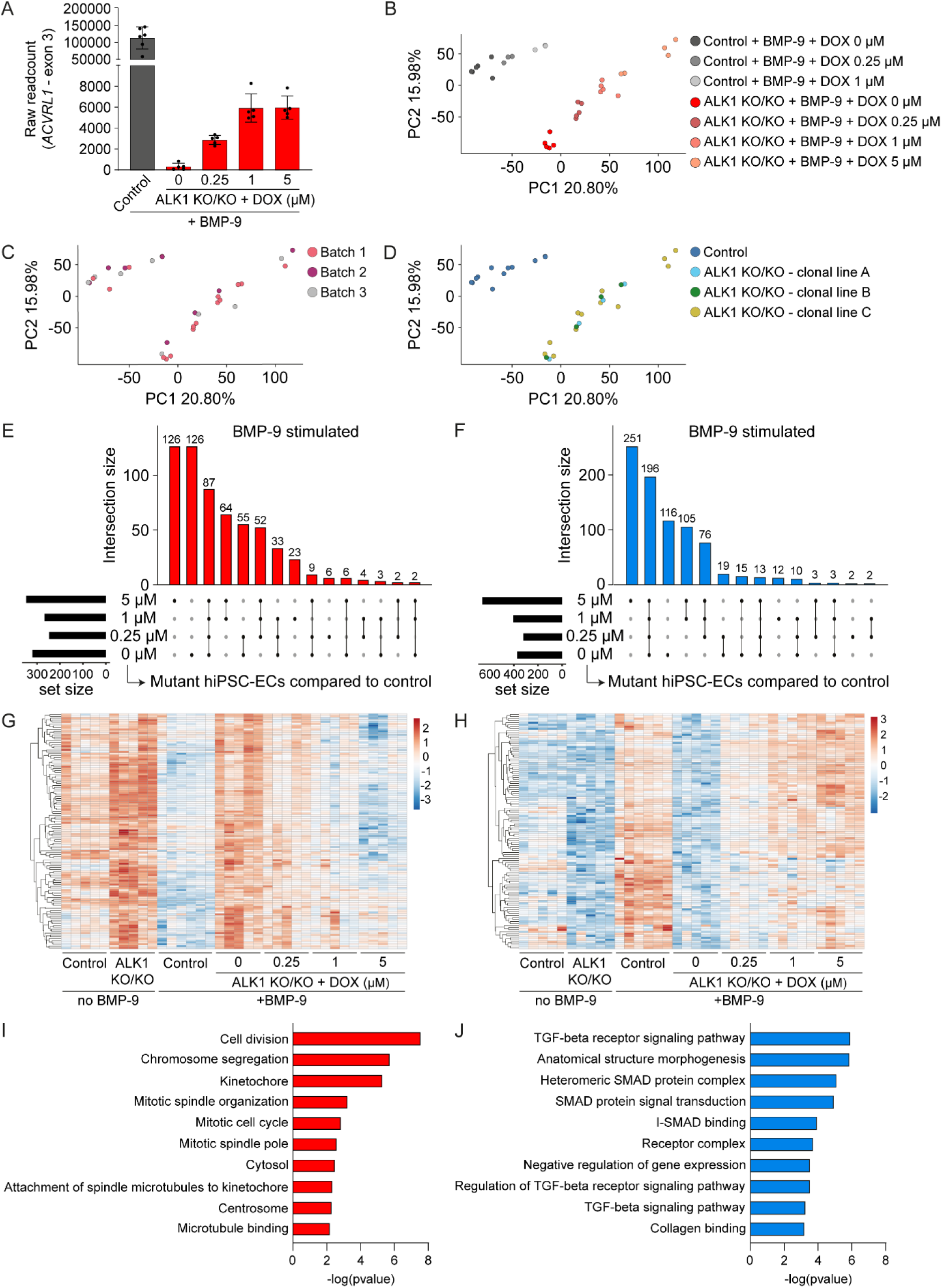
Doxycycline-induced ALK1 restores gene networks related to cell cycling and SMAD-signaling in a dose-dependent manner in ALK1 knockout hiPSC-ECs. (**A-J**) Transcriptomic analysis of untreated and doxycycline-treated (72 hours) control and ALK1 knockout hiPSC-ECs after DMSO or 1 ng/mL BMP-9 stimulation for 12 hours. N= 3 (independent hiPSC-EC batches); n= 1-2 (one to two technical replicates per batch). (**A**) Quantification of raw read counts that match with the deleted region in exon 3 of *ACVRL1* in ALK1 knockout ECs. Reads observed in doxycycline-treated ALK1 knockout ECs originate from the *AAVS1* locus. (**B-D**) Principal-component (PC) plots depicting the experimental conditions (**B**), batch-origin (**C**) and genetic-origin (**D**) of the samples. (**E-F**) Upset plot depicting the size and overlap of up-(**E**) and downregulated (**F**) genes (logCPM > 1, log2FC > 1, and p-adj < 0.05) between the indicated comparisons. (**G-H**) Semi-supervised clustering using the up-(**E**) and downregulated (**F**) genes unique to the ALK1 knockout ECs (+BMP-9, 0 μM DOX) vs. control hiPSC-ECs (+BMP-9, 0 μM DOX) comparison. (**I-J**) Gene ontology analysis on the up-and downregulated genes identified in (**G-H**). DOX, doxycycline.

### ALK1 knockout hiPSC-ECs exhibit high proliferative activity

Partial deficiency of ALK1 in ECs has been previously associated with vascular hyperplasia in murine models and skin biopsies from patients with HHT (Alsina-Sanchís et al., 2018; Chen et al., 2014). To investigate if ALK1 KO/KO hiPSC-ECs show increased proliferation rates, we determined the proliferation rates in both control and KO/KO hiPSC-ECs, either unstimulated or stimulated with BMP-9. Immunofluorescent analysis followed by quantification of 5-ethynyl-2’-deoxyuridine (EdU) positive nuclei indicated that ALK1 KO/KO hiPSC-ECs had elevated proliferation rates under baseline conditions compared to control cells (Figures S3A and S3B). To determine if reintroducing ALK1 restores BMP-9 sensitivity in knockout cells, ALK1 KO/KO hiPSC-ECs were exposed to different concentrations of doxycycline for 72 hours, with BMP-9 (1 ng/mL) stimulation during the last 24 hours. Analysis of EdU-positive nuclei revealed a dose-dependent reduction in proliferation as doxycycline-induced ALK1 expression increased (Figures 4A and 4B). Figures 4B and S3B show that BMP-9 stimulation reduced proliferation in control hiPSC-ECs, whereas ALK1 KO/KO ECs remained unresponsive, indicating that BMP-9 fails to activate ALK1-dependent pathways required for cell cycle suppression, corroborating our transcriptional data (Figure 3G and 3I). To more precisely characterize the state of the cell cycle, we utilized curated gene sets that mark the G1/S, G2/M, and G1/G0 phases of the cell cycle (Tirosh et al., 2016). In the absence of BMP-9, we observed increased expression of the G1/S and G2/M gene sets in ALK1 KO/KO hiPSC-ECs, which remained high following BMP-9 stimulation (Figure 4C). Induction of ALK1 in the knockout cells through doxycycline treatment restored the expression levels of the G1/S and G2/M gene sets comparable to control cells. The expression levels of G1/G0-related genes were comparable across all subgroups (Figure 4C). Additionally, doxycycline-treated control cells did not show a change in expression of the cell cycle gene signatures (Figures S3C and S3D). These data corroborate earlier studies showing that the absence of ALK1 leads to increased proliferation rates in ECs, and that BMP-9 functions as a ligand promoting vascular quiescence (Bidart et al., 2012; David et al., 2008).

**Figure 4.**
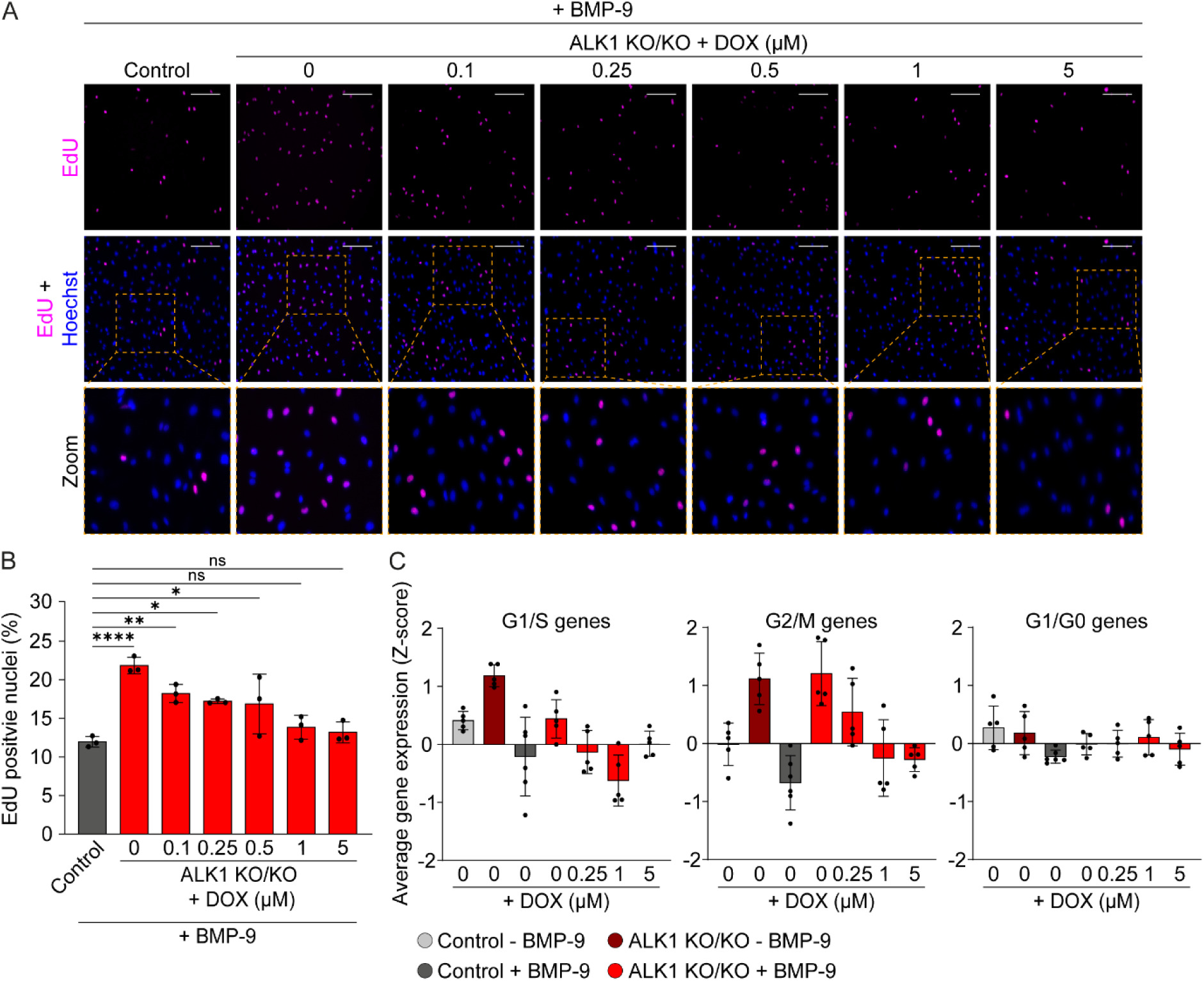
ALK1 knockout hiPSC-ECs exhibit high proliferative activity and impaired downregulation of G2/M-associated genes upon BMP-9 stimulation. (**A-B**) EdU analysis in control and ALK1 knockout hiPSC-ECs stimulated with 1 ng/mL BMP-9 for 24 hours. Knockout ECs were treated with the indicated concentrations of doxycycline for a total of 72 hours. (**A**) Representative immunofluorescence images showing EdU (magenta) and Hoechst (blue). Scale bars= 100 μm. (**B**) Quantification of EdU positive nuclei. N= 3 (independent hiPSC-EC batches); n= 9–18 (three to six images per experimental group per batch). (**C**) Average Z-score expression levels of curated gene sets associated with the G1/S, G2/M, and G0/G1 cell cycle phases in control and knockout hiPSC-ECs treated with doxycycline for 72h, either unstimulated or stimulated with 1 ng/mL BMP-9 for 12 hours. Data plotted as mean ±SD. Ordinary one-way ANOVA with Dunnett’s multiple comparisons test (**B**). * p < 0.05; ** p < 0.01; **** p < 0.0001; ns, not significant. DOX, doxycycline.

### Doxycycline-induced ALK1 restores tip/stalk cell specification defects in ALK1-deficient ECs

Mutant ALK1 mouse models of HHT have been reported to display impaired tip/stalk cell specification in ECs, contributing to excessive sprouting and formation of AVMs (Larrivee et al., 2012; Schmid et al., 2023). To investigate whether tip/stalk cell specification defects occurred in ALK1 KO/KO hiPSC-ECs, we cultured either knockout or isogenic control (ALK1 WT) hiPSC-ECs, both expressing Tdtomato, alongside genetically distinct control hiPSC-ECs (H2B-GFP; depicted in cyan) on Cytodex® 3 microcarrier beads, which were subsequently embedded in a fibrin gel to facilitate EC sprouting (Figure 5A). Both ALK1 WT and ALK1 KO/KO hiPSC-ECs contributed to sprouting events, as evidenced by immunofluorescence images (Figure 5B). Quantification of the Tdtomato-and GFP-positive nuclei located at the tip cell position revealed a strong bias, where ALK1 KO/KO cells were more frequently identified as tip cells in contrast to ALK1 WT cells (Figure 5C). This observation was consistent across varying experimental batches (Figure S4A). Doxycycline-induced expression of ALK1 restored this imbalance in a dose-dependent manner, and was associated with a reduced propensity of doxycycline-treated ALK1 KO/KO cells to generate sprouts (Figure 5D). Additionally, both ALK1 WT and ALK1 knockout hiPSC-ECs produced sprouts with similar length (Figure 5E). Importantly, treatment of the unrelated control hiPSC-ECs (H2B-GFP) with doxycycline did not affect the number or length of sprouts (Figures 5D and 5E; blue bars), indicating that doxycycline alone does not modulate sprouting behavior. Next, we assessed the expression pattern of genes that are typically enriched in endothelial tip cells (del Toro et al., 2010; Miyamura et al., 2024; Strasser et al., 2010; Yana et al., 2007), and found angiopoietin-2 (*ANGPT2*), apelin (*APLN*), C-X-C chemokine receptor type 4 (*CXCR4*), and matrix metalloproteinase-14 (*MMP14*) to be upregulated in unstimulated ALK1 KO/KO cells compared to control cells (Figures 5F and 5G). Upon BMP-9 stimulation, the expression levels of *APLN* and *CXCR4* were downregulated in control ECs, whereas expression levels remained elevated in knockout cells. Doxycycline-induced ALK1 in BMP-9 stimulated knockout ECs dose-dependently restored the expression abnormalities observed for *ANGPT2*, *CXCR4*, and *MMP14*, whereas *APLN* was dose-dependently downregulated but did not reach the expression levels observed in control cells. In addition, doxycycline treatment did not affect expression levels of *ANGPT2*, *APLN*, *CXCR4*, and *MMP14* in unstimulated and BMP-9 stimulated control hiPSC-ECs (Figures S4B and S4C). In summary, ALK1 KO/KO hiPSC-ECs show disrupted tip/stalk cell specification, increased tendency to sprout, and elevated expression of tip cell-enriched genes, all of which can be reversed upon doxycycline-induced ALK1 expression.

**Figure 5.**
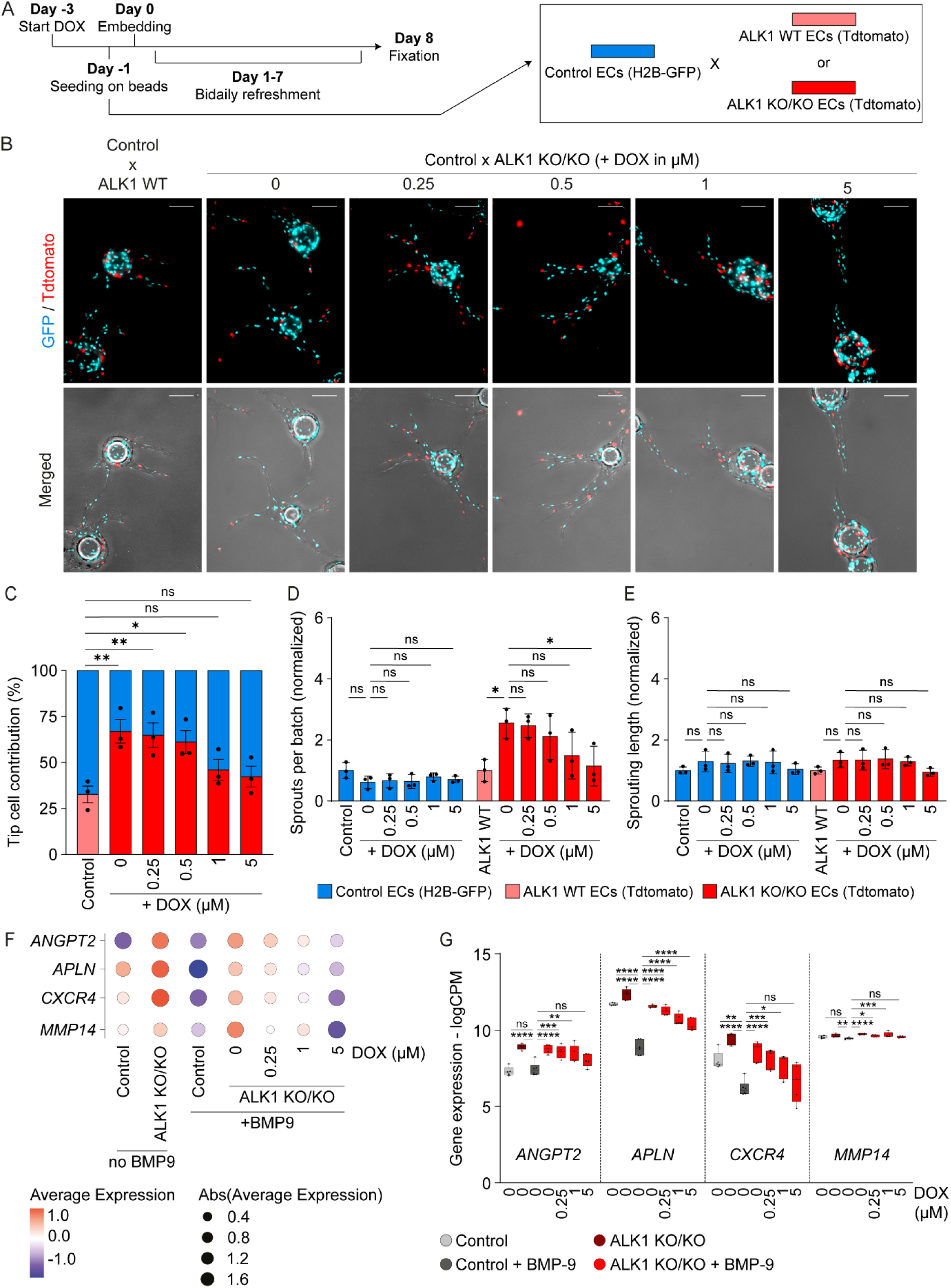
Endothelial tip/stalk cell selection defects are progressively restored upon doxycycline-mediated ALK1 induction. (**A**) Schematic depicting the competitive fibrin beads assay procedure. Control H2B-GFP hiPSC-ECs are depicted in blue, and ALK1 knockout ECs and the isogenic control (ALK1 WT) in red. (**B**) Representative immunofluorescence images for the indicated conditions. Scalebar= 50 μm. (**C**) Percentage of GFP+ and Tdtomato+ ECs located at the tip cell position. N= 3 (three independent batches of hiPSC-ECs); n= 10-20 (10 to 20 images analyzed for each experimental condition per batch). (**D-E**) Normalized number (**D**) and length (**E**) of sprouts per batch, guided by either a GFP+ or Tdtomato+ tip cell. N= 3 (three independent batches of hiPSC-ECs). (**F-G**) Expression levels of tip cell-associated genes depicted as a dotplot (**F**) and floating bar chart (**G**). Data plotted as mean ±SD. Ordinary one-way ANOVA with Dunnett’s (**C** and **G**) or Tukey’s multiple comparisons test (**D** and **E**). * p < 0.05; ** p < 0.01; *** p < 0.001; **** p < 0.0001; ns, not significant. DOX, doxycycline; logCPM, log counts per million.

### Doxycycline does not affect matrix metalloproteinase activity in hiPSC-ECs

Thus far, we have demonstrated that doxycycline-mediated induction of ALK1 in knockout ECs restores cellular function in a dose-dependent manner, including disrupted TGF-β/BMP-signaling pathways, excessive cellular proliferation, and abnormal tip/stalk cell specification. However, it is widely recognized that doxycycline itself suppresses the production and function of matrix metalloproteinases (MMPs) in ECs, thereby reducing extracellular matrix degradation and subsequently impacting both angiogenesis and vascular permeability (Formica et al., 2025; Hanemaaijer et al., 1998; Kim et al., 2005; Su et al., 2013; Wiggins-Dohlvik et al., 2016). To confirm that the beneficial outcomes observed in our model are caused by doxycycline-induced ALK1 rather than the doxycycline treatment itself, we first assessed the gene expression profiles of MMPs relevant to EC physiology (Figures S5A and S5B). *MMP1*, *MMP3*, *MMP9*, and *MMP13* were expressed at low levels (logCPM < 1) in both unstimulated and BMP-9 stimulated control and knockout hiPSC-ECs, whereas *MMP2*, *MMP10*, and *MMP14* displayed robust expression. Increasing concentrations of doxycycline progressively increased the expression of *MMP10*, whereas *MMP2* and *MMP14* displayed dose-dependent downregulation. Subsequently, we quantified the enzymatic activity of MMP-2 and MMP-14 in both untreated and doxycycline-treated control and knockout hiPSC-ECs. Although the overall levels of MMP-2 were reduced in control and knockout cells following exposure to 5 µM of doxycycline, no significant changes in active MMP-2 was observed (Figures S5C and S5D). Interestingly, despite similar total MMP-2 levels in both control and knockout cell populations, the fraction of active MMP-2 was elevated in ALK1 KO/KO hiPSC-ECs, which may suggest an increased ability to degrade the extracellular matrix. Furthermore, the levels of active MMP-14 were similar between control and knockout hiPSC-ECs and showed no changes in response to doxycycline treatment (Figure S5E). In addition, as previously observed, doxycycline had no effect on the number or length of sprouts formed and guided by control H2B-GFP hiPSC-ECs in the three-dimensional sprouting assay (Figures 5D and 5E; blue bars). Together, these data support the observed rescue effects being primarily due to the induction of ALK1 rather than unintended effects of doxycycline itself.

## DISCUSSION

In this study, we created a tunable human *in vitro* platform to investigate the dose-dependent effects of ALK1 on EC function, with direct implications for understanding the vascular pathology underlying ALK1-related HHT.

We first generated *ACVRL1* knockout hiPSCs and reintroduced wild-type *ACVRL1* in the *AAVS1* locus under a doxycycline-inducible system. This enabled us to titrate ALK1 expression levels within a certain range, allowing us to model the threshold-dependent effect of ALK1 signaling in ECs. Complete loss of ALK1 disrupts BMP-9 induced SMAD1/5 signaling, thereby facilitating uncontrolled EC proliferation. Reactivation of ALK1 expression, even at low levels, was sufficient to restore responsiveness towards BMP-9 and revert the proliferation defects. Transcriptomic analysis further supported these observations, revealing that ALK1-deficient hiPSC-ECs fail to control gene programs typically regulated by BMP-9, particularly those involved in cell cycling and extracellular matrix remodeling. ALK1 reinduction restored these gene networks in a dose-dependent manner, including the repression of mitotic and proliferative gene sets and the activation of genes linked to TGF-β/BMP pathway signaling. This is consistent with prior reports linking ALK1 deficiency to cell proliferation defects, possibly due to increased sensitivity towards VEGF (David et al., 2007; Kerr et al., 2015; Scharpfenecker et al., 2007; Shao et al., 2009; Thalgott et al., 2018; Upton et al., 2009).

In agreement with previous reports, ALK1-deficient hiPSC-ECs displayed altered tip/stalk cell specification dynamics, favoring a tip cell phenotype and producing more sprouts (Ahmed et al., 2023; Kerr et al., 2015; Larrivee et al., 2012; Schmid et al., 2023). Reinduction of ALK1 expression effectively reversed these phenotypic abnormalities, highlighting its critical role in EC fate determination. The transcriptional landscape of knockout cells reflected elevated expression of tip cell-enriched genes such as *ANGPT2*, *CXCR4*, and *APLN*, which were insensitive to BMP-9 in the absence of ALK1. The restoration of their expression upon doxycycline-induced ALK1 further underscores the specificity and importance of ALK1 signaling in regulating angiogenic sprouting dynamics.

Our model provides a possible explanation for tissue-specific vulnerability in HHT in humans. The dose-dependent response to ALK1 reactivation may suggest that vascular beds with inherently low *ACVRL1* expression are more prone to develop AVMs upon partial gene loss. This hypothesis parallels findings in ENG-related HHT, where regions with low *ENG* levels displayed increased susceptibility to lesion formation (Galaris et al., 2021). Experimentally assessing baseline *ACVRL1* levels across vascular beds will be essential to benchmark our model. At the same time, the ability to precisely titrate ALK1 in human ECs provides a relevant platform to investigate how basal gene dosage influences regional lesion development.

Given that doxycycline is known to modulate MMP activity and influence angiogenesis, we performed control experiments to exclude potential side effects of doxycycline treatment (Formica et al., 2025; Hanemaaijer et al., 1998; Kim et al., 2005; Su et al., 2013; Wiggins-Dohlvik et al., 2016). We observed modest changes in *MMP* gene expression in response to doxycycline, including a dose-dependent increase in *MMP10* and decrease in *MMP2* and *MMP14* transcripts. Functionally, no changes in active MMP-2 or MMP-14 levels were observed, nor did doxycycline treatment affect sprouting behavior in control cells. In contrast, levels of active MMP-2 were elevated in ALK1-deficient ECs, indicating increased extracellular matrix remodeling activity which could potentially contribute to HHT pathogenesis (Nagalingam et al., 2018; Xu et al., 2004). The doxycycline concentrations (0–5 µM) used in this study are substantially lower than those reported to inhibit MMP activity in previous studies. In the context of EC biology, Hanemaaijer et al. demonstrated that 10–50 µM doxycycline inhibits active MMP-8 and MMP-9, but not MMP-2 (Hanemaaijer et al., 1998). Similarly, Formica et al. reported a modest reduction in total MMP-2 levels at ∼20 µM, with the active MMP-2 pool remaining unaffected, whereas complete inhibition of both MMP-2 and MMP-9 was observed at ∼100 µM (Formica et al., 2025). Some studies have shown that doxycycline concentrations of ∼20–40 µM are required to suppress MMP-2 and MMP-9 activity (Kim et al., 2005; Su et al., 2013), whereas another study found no effect on active MMP-9 at 20 µM in rat lung microvascular ECs (Wiggins-Dohlvik et al., 2016). In a clinical setting, treatment with doxycycline at a dose of 100 mg twice a day did not improve Epistaxis Severity Scores (ESS) in HHT patients (McWilliams et al., 2022; Thompson et al., 2022). Additionally, Thompson et al. reported no reduction in circulating MMP-9 levels following treatment. Together, our observations support that the phenotypic rescue observed in ALK1-deficient cells arises from reinduced ALK1 expression itself, rather than from doxycycline-mediated MMP suppression.

Though the induced expression level of ALK1 achieved in our system (∼10% compared to control hiPSC-ECs) was sufficient to recapitulate key aspects of HHT pathology within the experimental frameworks utilized, this limited induction efficiency may represent a constraint when extending the approach to other genes or diseases. Several factors could underlie this submaximal expression, including monoallelic integration of the inducible construct, suboptimal promoter and regulatory element configurations within the DNA cassette (Blanch-Asensio, Ploessl, et al., 2024; Johnstone et al., 2025), or locus-specific effects associated with *AAVS1* targeting. Indeed, studies have suggested that other safe harbor loci, such as citrate lyase beta-like (*CLYBL*), may improve the expression profiles of transgenes (Blanch-Asensio et al., 2023; Cerbini et al., 2015; Guichardaz et al., 2024). In addition, gene-intrinsic differences in transcription and translation may further affect the efficacy of inducible systems.

Although our tunable *in vitro* platform offers important insights into the dose-dependent effects of ALK1 on endothelial function, it represents a simplified system that cannot fully mimic the complex 3D architecture, biomechanical forces, and cellular diversity found *in vivo*. Advanced systems such as vessel-on-chip (VoC) platforms, which incorporate microfluidic flow, three-dimensional extracellular matrix components, and multicellular interactions, provide a more physiologically relevant environment to study how ALK1 dosage influences endothelial behavior under dynamic shear stress and within the vascular niche (Fang et al., 2024; Orlova et al., 2022; Soon et al., 2022; Vila Cuenca et al., 2021). Indeed, recent work has shown that increased flow conditions and vessel enlargement, hallmarks of AVMs, emerge only in these more complex models (Fang et al., 2024). Moreover, the coexistence of knockout and wild-type ECs and the resulting non-cell-autonomous effects on lesion formation emphasize the necessity of spatial and cellular heterogeneity. Furthermore, recent advances have enabled the incorporation of immune cells into VoC systems and improved the long-term stability of engineered vasculature, preventing regression during extended culture periods (Floryan et al., 2025; Jäger et al., 2025; van Dijk et al., 2020). These models more accurately mimic tissue-specific microenvironments, enabling investigation of how ALK1-related signaling integrates with flow-mediated cues, perivascular cell interactions, and immune components to drive regional susceptibility to vascular lesion formation. Integrating our doxycycline-inducible ALK1 system with VoC technology would therefore provide a powerful approach to dissect the biological and microenvironmental factors contributing to HHT pathogenesis, bridging the gap between simplified cellular assays and complex *in vivo* conditions.

Our data show the importance of maintaining ALK1 expression above a functional threshold to ensure endothelial cell integrity, responsiveness to BMP-9, and angiogenic fate. The dose-dependent rescue of endothelial dysfunction sets the stage for understanding the tissue-specific manifestations of ALK1-related HHT, where regions of low basal *ACVRL1* expression may be more susceptible to AVM formation upon genetic inactivation. Future work will focus on combining this tunable humanized platform with VoC methodologies to further understand HHT pathobiology and investigate potential therapeutic strategies aimed at restoring ALK1 activity.

## EXPERIMENTAL PROCEDURES

### CRISPR-Cas9-mediated deletion of ALK1

Gene editing of LU99_*AAVS1*-bxb-v2 (LUMCi004-A-1) was performed using the NEON Transfection System (Invitrogen) with the following electroporation settings: 1200 V, 30 ms, 1 pulse. A total of 1.0 x 10^5^ cells were electroporated with a Cas9-RNP complex targeting exon 3 of *ACVRL1*. The Cas9-RNP complex was assembled by incubating Cas9 protein (Cas9 nuclease V3, IDT) with a multiplex mixture of three sgRNAs (Gene Knockout Kit v2 - human - *ACVRL1*, Synthego), following the manufacturer’s protocol. Briefly, sgRNAs were dissolved to a final concentration of 100 µM (100 pmol/µl) and mixed with Cas9 protein at a 2:1 molar ratio. After electroporation cells were plated on Corning® Synthemax® II-SC substrate (Merck, # CLS3535-1EA) in mTeSR Plus Medium (STEMCELL Technologies, #100-0276), supplemented with CloneR™ 2 (STEMCELL Technologies, #100-0691). Following recovery, 1000 cells were plated on a 10cm Synthemax II-SC-coated dish in mTeSR Plus with CloneR™ 2. Single cell-derived colonies were manually picked and screened for the *ALK1* knockout genotype. Genomic DNA was isolated using QuickExtract™ DNA Extraction Solution (Lucigen, #QE0905T), and the targeted region amplified by PCR (Terra PCR Direct Polymerase Mix, TaKaRa). PCR products were analyzed by gel electrophoresis, and candidate *ALK1*-KO/KO clones validated by Sanger sequencing (Leiden Genome Technology Centre using the ABI3730xl system). The sgRNA target sequences and primer pairs used for genotyping are listed in Table S1.

### Construction of doxycycline-inducible ACVRL1 donor plasmid

The doxycycline-inducible *ACVRL1* donor vector was generated by two sequential cloning steps using the NEBuilder HiFi DNA Assembly kit (NEB). All assembly reactions were performed for one hour at 50 °C, following the manufacturer’s protocol. In the first step, a generic doxycycline-inducible donor plasmid was assembled. A polyadenylation (polyA) sequence and a plasmid encoding the all-in-one Tet-On 3G inducible system (TRE-3G_lacZα-MCS_WPRE_pCAG_Tet-On^®^3G_T2A, Addgene #229783) were PCR-amplified and purified using the Wizard SV Gel and PCR Clean-Up System kit (Promega, #A9281). These fragments were inserted into the NheI-(NEB, R3131S) linearized pBR_attB(bxb)_lox donor plasmid (Addgene, #183762). In the second step, the final doxycycline-inducible *ACVRL1* donor vector was constructed by digesting the generic doxycycline-inducible donor plasmid with NotI and NheI (NEB, R3189S and R3131S, respectively), and inserting three DNA fragments. These fragments, each containing short homology arms, were obtained via PCR-amplification (Tdtomato, Addgene #198041), or ordered as gene fragments from Twist Bioscience (wildtype_*ACVRL1* and 3xNLS). Primer sequences and synthetic gene fragment designs are provided in Tables S1 and S2, respectively.

### Targeted integration of the doxycycline-inducible ACVRL1 cassette using STRAIGHT-IN

The integration of the doxycycline-inducible *ACVRL1* donor plasmid into the *AAVS1* locus of *ACVRL1*-null hiPSCs was performed as previously described (Blanch-Asensio, Grandela, et al., 2024). Briefly, the donor plasmid and a Bxb1 recombinase-expressing plasmid (Addgene #51271) were transfected into *AAVS1*-bxb-v2 *ACVRL1*-null hiPSCs using Lipofectamine™ Stem reagent (ThermoFisher, STEM00008). Plasmid amounts were adjusted according to vector size. Following transfection, cells were passaged and selected with zeocin (ThermoFisher, R25001) to enrich for clones that had correctly integrated the cassette. Integration efficiency was assessed by ddPCR detecting *attP* and *attR* sites, as previously described (Blanch-Asensio, Grandela, et al., 2024). To excise the auxiliary sequences flanking the integrated cassette, hiPSCs were transfected with a Cre recombinase-expressing plasmid (Addgene #183812) and subsequently selected with puromycin (ThermoFisher, J67236-XF). Excision was confirmed by ddPCR quantifying the loss of *BleoR*. Primer and probes sequences used for integration and copy number validation are listed in Table S1.

### hiPSC cell culture and Genomic DNA Extraction

hiPSCs were cultured on Vitronectin XF™-coated plates (STEMCELL Technologies, #07180). Genomic DNA was extracted using DNeasy Blood & Tissue Kit (QIAGEN, #69504) performed according to the manufacturer’s instructions.

### Droplet Digital PCR

ddPCR was carried out using a thermocycler, the Q200 AutoDG and QX200 Droplet Digital PCR System, and analyzed with QuantaSoft software (all Bio-Rad) as previously described (Blanch-Asensio, Grandela, et al., 2024). Assays containing pre-mixed forward and reverse primers (each at 18 mM) along with a FAM-or HEX-labeled hydrolysis probe (5 mM) were either obtained from Bio-Rad (Roberts et al., 2017), or custom-designed following specific guidelines (Bell et al., 2018). Primer sequences and probes are listed in Table S1.

### Endothelial Cell Differentiation and Maintenance

Differentiation of hiPSCs into ECs, and subsequent enrichment using CD31 magnetic beads was performed using established protocols (Halaidych et al., 2018; Orlova, Drabsch, et al., 2014; Orlova, van den Hil, et al., 2014). Briefly, one day before initiating differentiation, hiPSCs were passaged on Vitronectin XF™-coated 6-well plates (STEMCELL Technologies, #07180) and maintained in StemFlex™ Medium (ThermoFisher, A3349401). Mesoderm induction was initiated by refreshing the medium with B(P)EL supplemented with 8.0 µM CHIR99021 (Tocris Bioscience, #4423). On days three, six, and nine, the medium was refreshed with B(P)EL supplemented with 50 ng/mL VEGF (Miltenyi Biotec, #130-109-386) and 10 µM SB431542 (Tocris Bioscience, #1614). ECs were isolated on day 10 using CD31-coated magnetic beads (Invitrogen, #11155D) and expanded for four days on 0.1% gelatin-coated (Sigma-Aldrich, G1890) flasks in endothelial cell complete growth medium (EC-CGM), composed of human endothelial serum-free medium (EC-SFM, Invitrogen, #11111-044) supplemented with 30 ng/mL VEGF (Miltenyi Biotec, #130-109-386), 20 ng/mL bFGF (Miltenyi Biotec, #130-093-842), and 1% human platelet-poor serum (hPPS, Sigma-Aldrich, P2918). ECs were frozen in cryopreservation medium; 50% FBS (Biowest, S1860), 40% EGM-2 (Lonza, CC-3162), and 10% DMSO (Sigma-Aldrich, D2650).

### Flow Cytometry

hiPSCs were maintained in StemFlex™ Medium (ThermoFisher, A3349401), whereas hiPSC-ECs were thawed on 0.1% gelatin-coated (Sigma-Aldrich, G1890) culture plastics and maintained in EC-CGM, either in the absence or presence of doxycycline (Sigma, D9891-1G). Cells were dissociated using TrypLE Select (Gibco, #12563029), and washed once in buffer (0.5% BSA (Biowest, #S1860), 5.0 mM 0.5 M EDTA (Sigma-Aldrich, E5134) in PBS (Gibco, #14190-094)). Staining with primary antibody was performed at room temperature for 30 minutes, followed by three washes with buffer. In case of non-conjugated primary antibodies, cells were stained with Alexa Fluor™ secondary antibodies at 1:1000 (ThermoFisher) for 15 minutes at room temperature in the dark, followed by three washes with buffer. Stained cells were then incubated with DAPI for five minutes (1:1000, Invitrogen, D3571), and analyzed on a MACSQuant® VYB Flow Cytometer (Miltenyi Biotec) with the following configuration: Blue/488 FITC, A488: 525/50; Yellow/561 PE: 586/15; APC: 661/20; APC-Cy7: 750LP. Collected data was exported and further analyzed using FlowJo software (BD, Version 10.8.0). Antibodies are listed in Table S3.

### Western blotting

hiPSC-ECs were thawed on 0.1% gelatin-coated (Sigma-Aldrich, G1890) culture plastics and maintained in EC-CGM for 66 hours, either in the absence or presence of doxycycline (Sigma, D9891-1G). Medium was then replaced with EC-SFM (Invitrogen, #11111-044) supplemented with 1% human platelet-poor serum (hPPS, Sigma-Aldrich, P2918) for a total of six hours. During the last 45 minutes, cells were either non-stimulated or stimulated with recombinant human BMP-9 protein (R&D systems, #3209-BP/CF) at the indicated concentrations. Culture plates were put on ice and washed once with ice cold PBS (Gibco, #14190-094). After aspiration, cells were lysed using RIPA (ThermoFisher, #89901) supplemented with cOmplete™ EDTA-free Protease Inhibitor Cocktail (Roche, #11873580001) and PhosSTOP™ (Roche, #4906845001). Protein concentrations were determined using the Micro BCA™ Protein Assay Kit (ThermoFisher, #23235) and analyzed on a SpectraMax Microplate Reader (Molecular Devices) according to manufacturer’s instructions. For Western blotting, 2.5-5.0 µg of protein lysate was used and processed using the Criterion™ Vertical electrophoresis and blotting system and reagents (all from Bio-Rad), following the general guidelines supplied by the manufacturer. Detection was performed using HRP-conjugated secondary antibodies and either the Clarity™ Western ECL Substrate (Bio-Rad, #1705061) or Clarity™ Max Western ECL Substrate (Bio-Rad, #1705062). Signal acquisition was conducted using a ChemiDoc Imaging System (Bio-Rad), and acquired band intensities were quantified using ImageJ. Antibody information is listed in Table S3.

### Immunofluorescence labeling

hiPSC-ECs were thawed on 0.1% gelatin-coated (Sigma-Aldrich, G1890) µ-Slide 8 well culture plastics (ibidi, #80821) and maintained in EC-CGM for 66 hours, either in the absence or presence of doxycycline (Sigma, D9891-1G). After, medium was replaced with EC-SFM (Invitrogen, #11111-044) supplemented with 1% human platelet-poor serum (hPPS, Sigma-Aldrich, P2918) for a total of six hours. During the last 45 minutes, cells were either non-stimulated or stimulated with 10 pg/mL recombinant human BMP-9 protein (R&D systems, #3209-BP/CF) at the indicated concentrations. Cells were fixed with 4% paraformaldehyde in PBS (ThermoFisher, J61899-AP) for 10 minutes at room temperature, permeabilized with 0.1% Triton X-100 in PBS (Sigma, X100-100ML), and stained with the indicated primary and secondary antibodies listed in Table S3. Images were acquired using an EVOS M7000 system (ThermoFisher) equipped with a 4x dry objective.

### Cell proliferation assay

hiPSC-ECs were thawed on 0.1% gelatin-coated (Sigma-Aldrich, G1890) culture plastics and maintained in EC-CGM for 48 hours, either in the absence or presence of doxycycline (Sigma, D9891-1G). After 48 hours, the medium was replaced with EC-SFM (Invitrogen, #11111-044) supplemented with 1% human platelet-poor serum (hPPS, Sigma-Aldrich, P2918), and with or without doxycycline and 1 ng/mL recombinant human BMP-9 protein (R&D systems, #3209-BP/CF. After 24 hours, cells were stained with the Click-iT™ Plus EdU Cell Proliferation Kit for Imaging (ThermoFisher, C10637) according to manufacturer’s instructions. EdU compound was added at a final concentration of 10 μM for 60 minutes. Samples were counterstained with DAPI (1:1000, Invitrogen, D3571), and imaged using an EVOS M7000 system (ThermoFisher) using a 4x dry objective.

### Bulk mRNA Sequencing

RNA was extracted using NucleoSpin™ RNA Columns (MN, #740955.50S), prepared with poly A enrichment, and sequenced on a NovaSeq X Plus Series instrument (PE150) at Novogene (Munich, Germany). Raw reads were pre-processed and aligned to the hg19 human reference genome using the bioWDL pipeline (Vorderman et al., 2021). Read counts were processed with edgeR as recommended by the authors, followed by a likelihood ratio test to identify differentially expressed genes (Robinson et al., 2010). Significance was defined as adjusted p-value < 0.05, logCPM > 1, and |logFC| > 1. All calculations were done using the R programming language version 4.5.0. The ggplot2 package (3.5.2) was used to generate dot-, PCA-, and volcano-plots, whereas heatmaps were created with pheatmap (1.0.13). The expression pattern of cell cycle-related genes was determined by applying z-score normalization to the expression values in counts per million of pre-selected gene sets (Table S4), and the average z-scores were then calculated per sample. Gene enrichment analysis was performed as follows, genes found to be differentially expressed were analyzed for functional enrichment using the DAVID tool (Sherman et al., 2022). The analysis was performed with default settings, using *Homo sapiens* as the background reference. Enrichment was assessed across Gene Ontology (GO) categories: molecular function (GO-MF), biological process (GO-BP), cellular component (GO-CC), and the KEGG pathway database.

### Fibrin Beads Assay

Preparation of reagents and cells was performed using an established protocol (Winters et al., 2016). Briefly, unrelated control H2B-GFP, ALK1 WT and ALK1 KO/KO hiPSC-ECs were seeded onto Cytodex 3 microcarrier beads (Merck, GE17-0485-01) at a 1:1 ratio, and embedded in a fibrin gel. After solidification of the fibrin gel, 20,000 human brain vascular pericytes (SanBio, SCC1200) were added to each well. Cultures were kept in EGM-2 supplemented with or without doxycycline (Sigma, D9891-1G) and refreshed bi-daily. After one week, cultures were fixed with 2% paraformaldehyde in PBS (ThermoFisher, J61899-AP) for 60 minutes at room temperature in the dark, permeabilized with blocking solution (1% BSA and 0.2% Triton-X100 in PBS) for two hours, and counterstained overnight with DAPI (1:1000). After washing three times with PBS, images were acquired using an EVOS M7000 system (ThermoFisher) equipped with a 10x dry objective.

### MMP-2 and MMP-14 activity assay

Activity levels of MMP-2 and MMP-14, and total MMP-2 levels were determined using the Human MMP-2 and MMP-14 activity assays following manufacturer’s instructions (QuickZyme Biosciences, QZBMMP-2Hv2 and QZBMMP-14H). Briefly, hiPSC-ECs were maintained for 72 hours in EC-CGM and processed as follows; for active MMP-14 levels, sample extracts were prepared using the provided extraction buffer, whereas supernatant of each culture was collected to determine active and total MMP-2. MMP-2 and MMP-14 were captured on antibody-coated plates, and activity was detected by measuring the enzymatic conversion of a chromogenic substrate. P-aminophenyl mercuric acetate was added to convert all inactive MMP-2 to its active form, followed by enzymatic conversion of the chromogenic substrate. Absorbance was measured on a SpectraMax Microplate Reader (Molecular Devices).

### Statistical Analysis

Data are plotted as mean ± SD. Statistical analyses were performed using PRISM (GraphPad Software, version 10.2.3). For comparisons between two groups, statistical significance was determined using a two-tailed unpaired Student’s t-test or a two-tailed Mann-Whitney test for non-normally distributed data. For comparisons involving more than two groups with one or two variables, an ordinary one-or two-way ANOVA was performed followed by a post hoc test, as described in each figure legend. Statistical significance was defined as p < 0.05, with significance levels indicated as follows: *p < 0.05, **p < 0.01, ***p < 0.001, ****p < 0.0001.

## ACKNOWLEDGEMENTS

The authors would like to thank Leon Mei from the Sequencing Analysis Support Core (SASC) of the Leiden University Medical Center (LUMC) for bioinformatics support and Franck Lebrin for discussion. This work was supported by the Netherlands Organ-on-Chip Initiative – NOCI (024.003.001), funded by the Ministry of Education, Culture and Science of the government of the Netherlands; and The Novo Nordisk Foundation Center for Stem Cell Medicine (NNF21CC0073729).

## AUTHOR CONTRIBUTIONS

S.J.v.K., designed, performed, and analyzed the research, performed bulk mRNA sequencing analyses, and wrote the manuscript; A.B.A., assisted with the design of the cloning and targeting strategy, and supplied required materials; J.L.G., performed and assisted with bioinformatics analyses; B.v.d.V., designed and performed the *ACVRL*-null hiPSCs experiments; C.M.A.H.F., designed the *ACVRL*-null hiPSCs experiments; R.P.D., designed the cloning and targeting strategy; C.L.M., conceptualized the study; V.V.O., conceptualized the study, and wrote the manuscript.

---

**Supplemental Figure 1.**
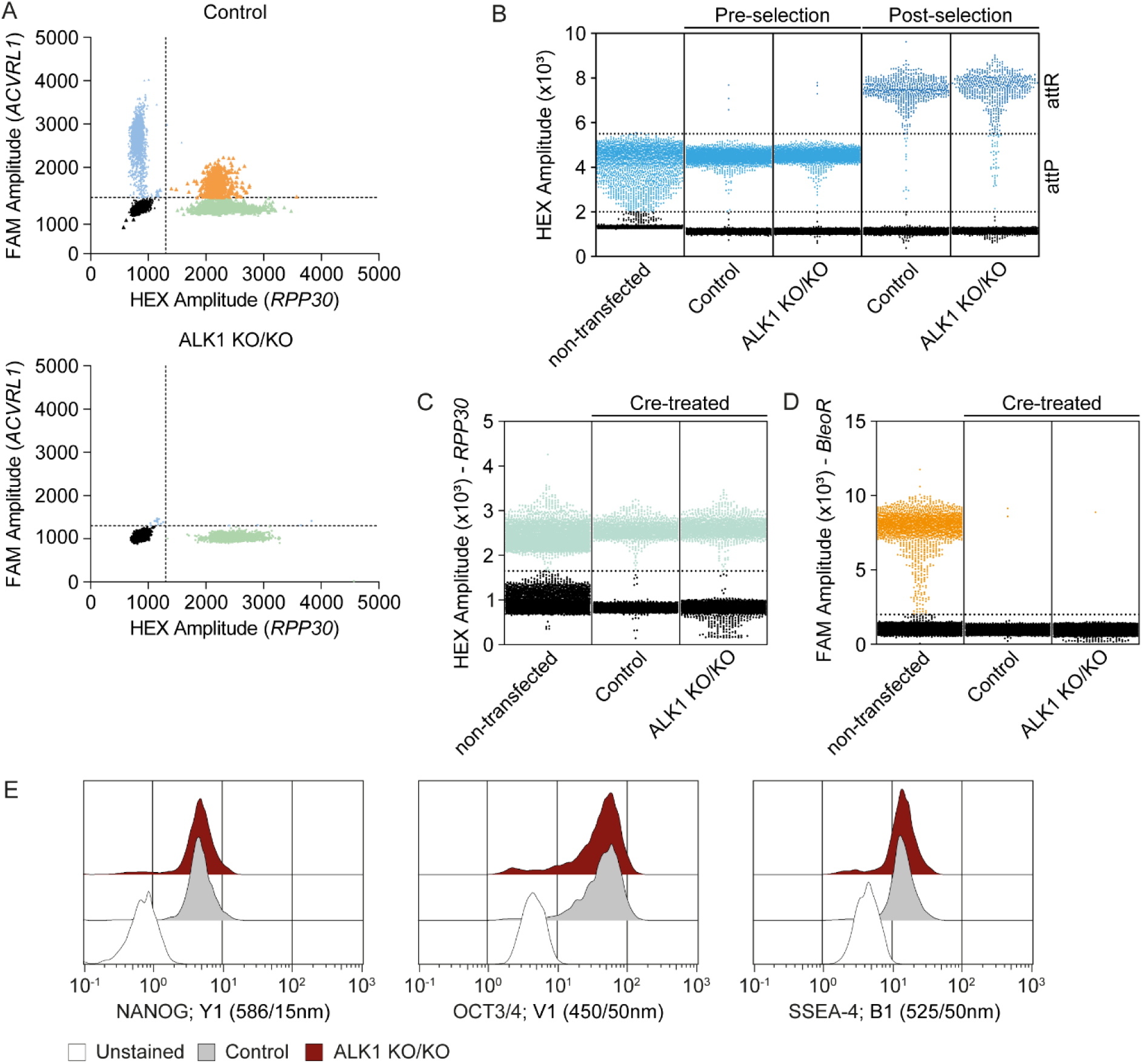
Related to Figure 1. Integration of a doxycycline-inducible ALK1 cassette into ALK1 knockout hiPSCs. (**A**) Representative ddPCR dot plots indicating the presence or absence of droplets positive for the *ACVRL1* and *RPP30* targeted sequences. (**B**) Representative ddPCR dot plots indicating the presence of droplets positive for *attP* (not integrated) or *attR* (integrated) sequences before and after zeocin selection. (**C-D**) Representative ddPCR dot plots indicating the presence of droplets positive for *RPP30* (**C**) or *BleoR* (**D**) sequences in Cre-treated hiPSCs. (**E**) Flow cytometry analysis of markers of the undifferentiated state NANOG, OCT3/4, and SSEA-4 in hiPSCs. *BleoR*, bleomycin.

**Supplemental Figure 2.**
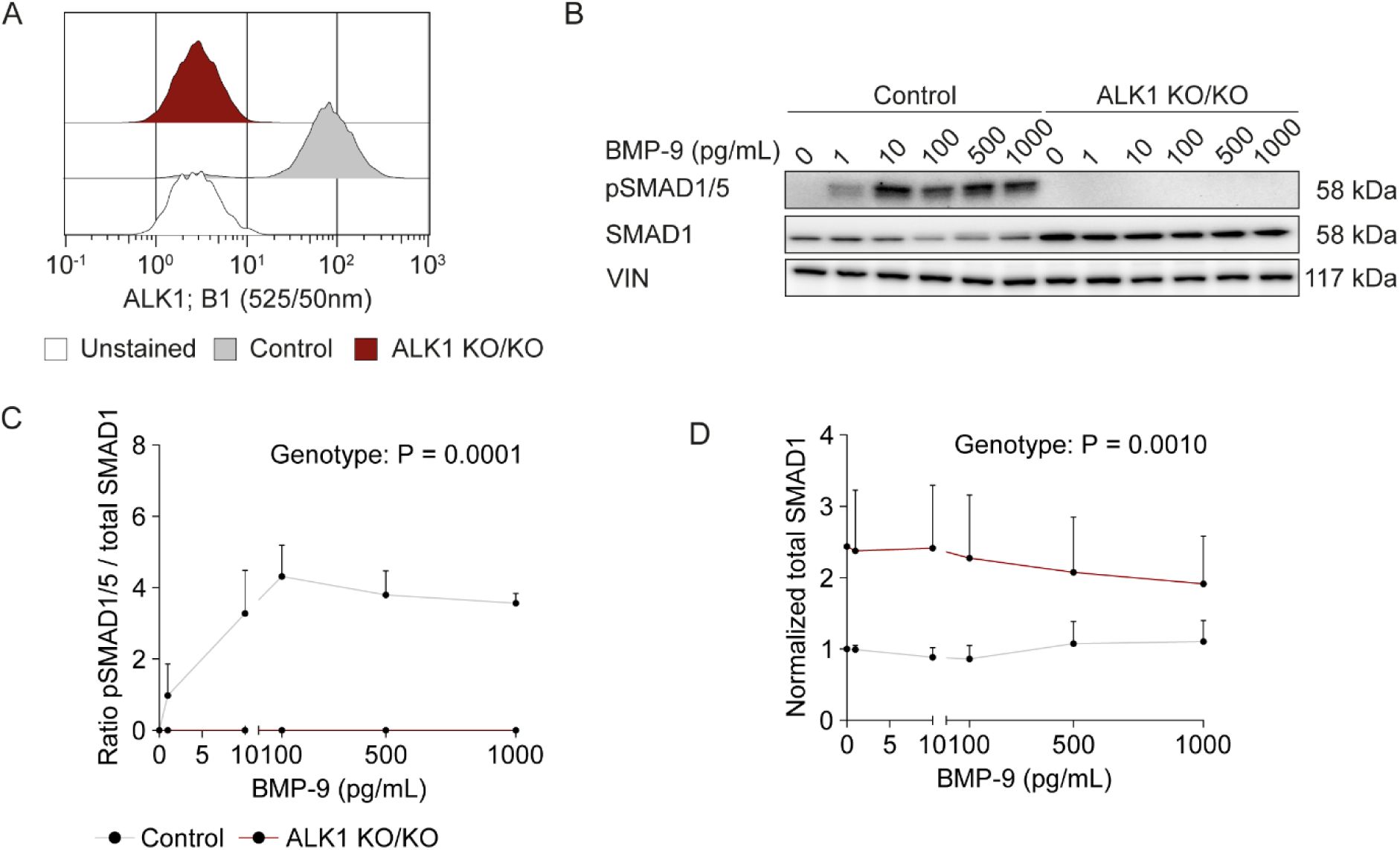
Related to Figure 1. ALK1 knockout hiPSC-ECs show altered SMAD1/5 activation dynamics upon BMP-9 stimulation. (**A**) Flow cytometry analysis of ALK1 in hiPSC-ECs. (**B-D**) Western blot analysis of extracts from serum starved control and ALK1 knockout hiPSC-ECs stimulated for 45min with the indicated BMP-9 concentrations. N= 3 (independent batches of hiPSC-ECs). (**B**) Representative Western blots for pSMAD1/5, total SMAD1, and Vinculin (VIN). (**C**) Quantification of pSMAD1/5 over total SMAD1 ratios. (**D**) Quantification of total SMAD1 protein levels normalized to VIN. Data plotted as mean ±SD. Ordinary two-way ANOVA test (**C,D**).

**Supplemental Figure 3.**
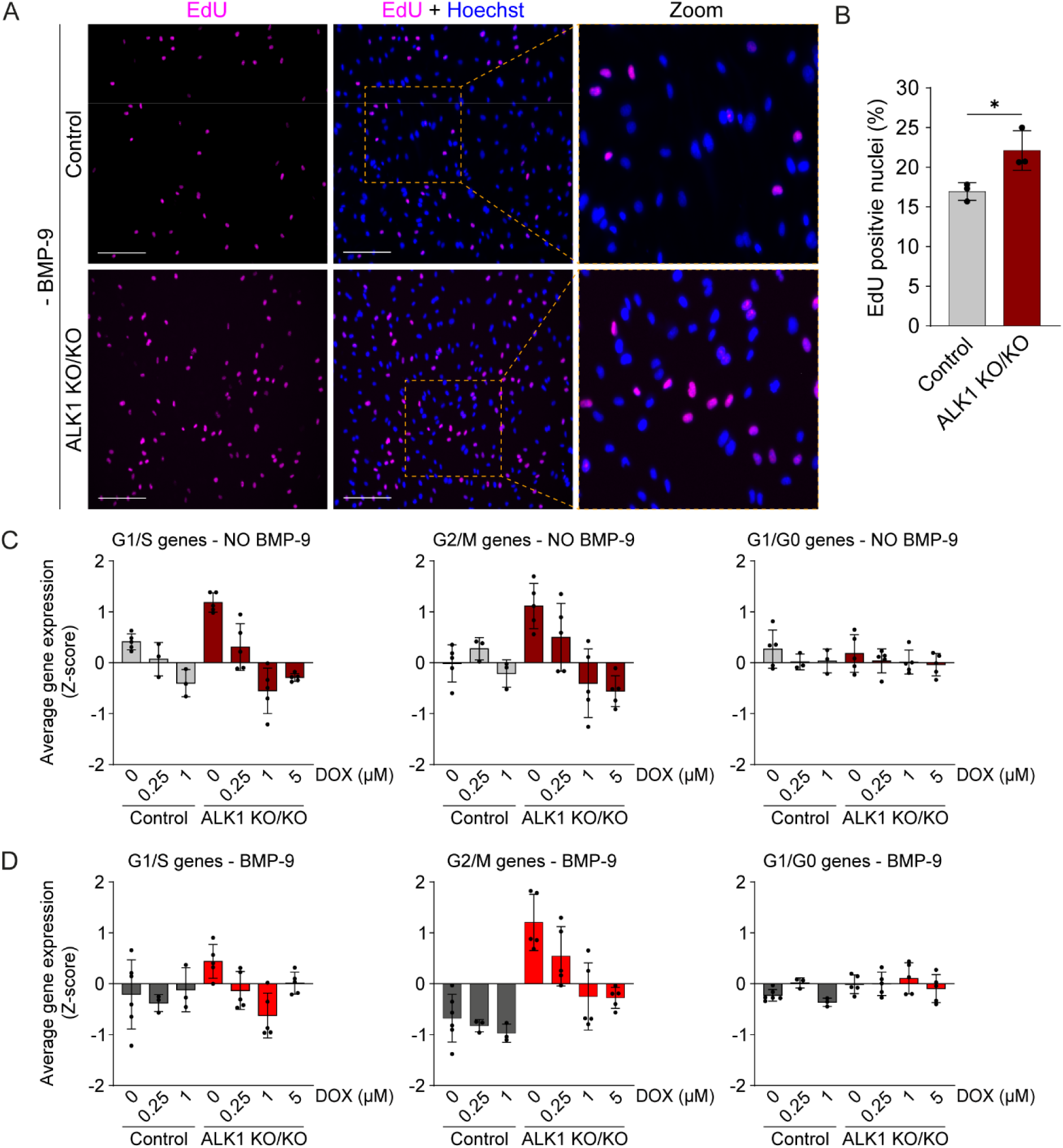
Related to Figure 4. ALK1 knockout hiPSC-ECs exhibit increased cellular proliferation under baseline conditions. (**A-B**) EdU analysis in control and ALK1 knockout hiPSC-ECs in the absence of BMP-9. (**A**) Representative immunofluorescence images showing EdU (magenta) and Hoechst (blue). Scale bars= 100 μm. (**B**) Quantification of EdU positive nuclei. N= 3 (independent hiPSC-EC batches); n= 9–18 (three to six images per experimental group per batch). (**C-D**) Average Z-score expression levels of curated gene sets associated with the G1/S, G2/M, and G0/G1 cell cycle phases in control and knockout hiPSC-ECs treated with doxycycline for 72h, either unstimulated (**C**) or stimulated (**D**) with 1 ng/mL BMP-9 for 12 hours. Data plotted as mean ±SDs. Two-tailed unpaired Student’s t-test (**B**). * p < 0.05. DOX, doxycycline.

**Supplemental Figure 4.**
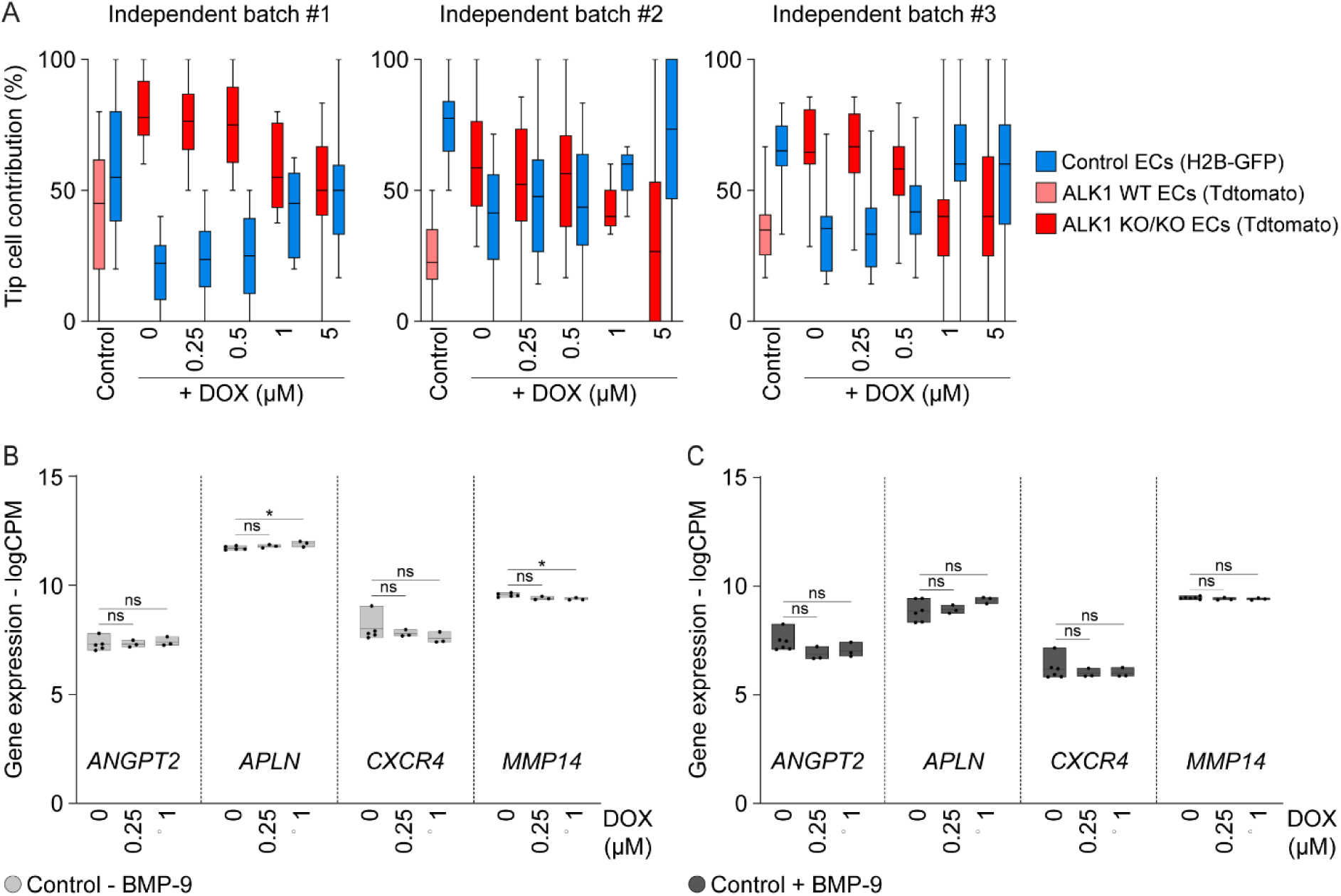
Related to Figure 5. Competitive fibrin beads assay quantification. (**A**) Percentage of GFP+ and Tdtomato+ ECs located at the tip cell position for each independent batch plotted in **Figure 5C**. (**B-C**) Expression levels of tip cell-associated genes in untreated and doxycycline-treated (72 hours) control hiPSC-ECs following 12 hours stimulation without (**B**) or with (**C**) 1 ng/mL BMP-9. N= 3 (independent hiPSC-EC batches); n= 1-2 (one to two technical replicates per batch). Data plotted as mean ±SD. Ordinary one-way ANOVA with Dunnett’s multiple comparisons test (**B**-**C**). * p < 0.05; ns, not significant. DOX, doxycycline; logCPM, log counts per million.

**Supplemental Figure 5.**
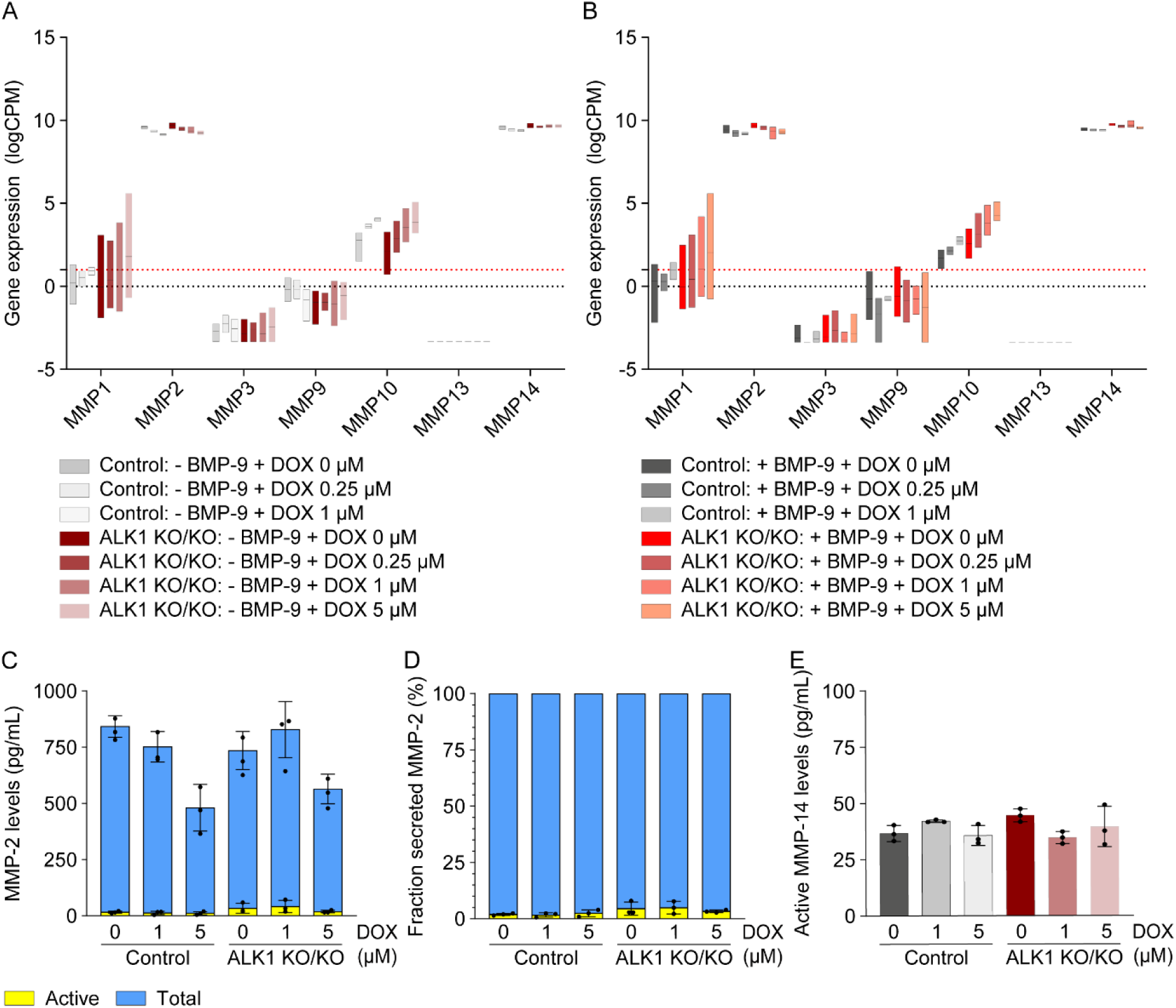
Doxycycline-treated hiPSC-ECs exhibit transcriptional changes but without functional alterations in MMP-related cell biology. (**A-B**) *MMP* expression levels in untreated and doxycycline-treated (72 hours) control and ALK1 knockout hiPSC-ECs following 12 hours stimulation with DMSO (**A**) or 1 ng/mL BMP-9 (**B**). N= 3 (independent hiPSC-EC batches); n= 1-2 (one to two technical replicates per batch). Black and red dotted lines marks logCPM values of 0 and 1, respectively. (**C**-**D**) Levels of active and total MMP-2 in doxycycline-treated (72 hours) control and ALK1 knockout hiPSC-ECs, either plotted as concentration (**C**) or as fraction (**D**). (**E**) Levels of active MMP-14 in extracts collected from doxycycline-treated (72 hours) control and ALK1 knockout hiPSC-ECs. Data plotted as mean ±SD. DOX, doxycycline; logCPM, log counts per million.

**Table S1:**
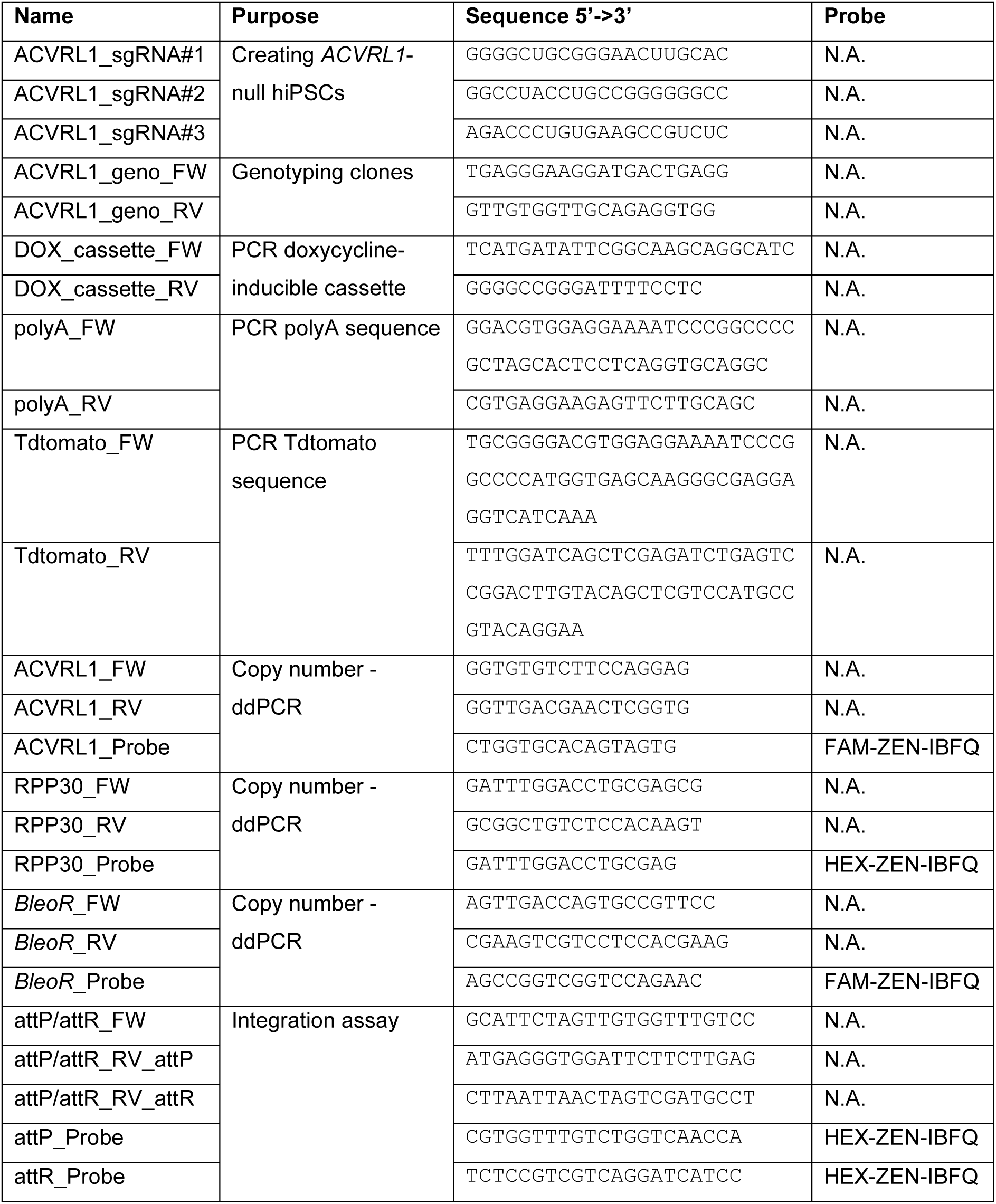
Primer and probe sequences.

**Table S2:**
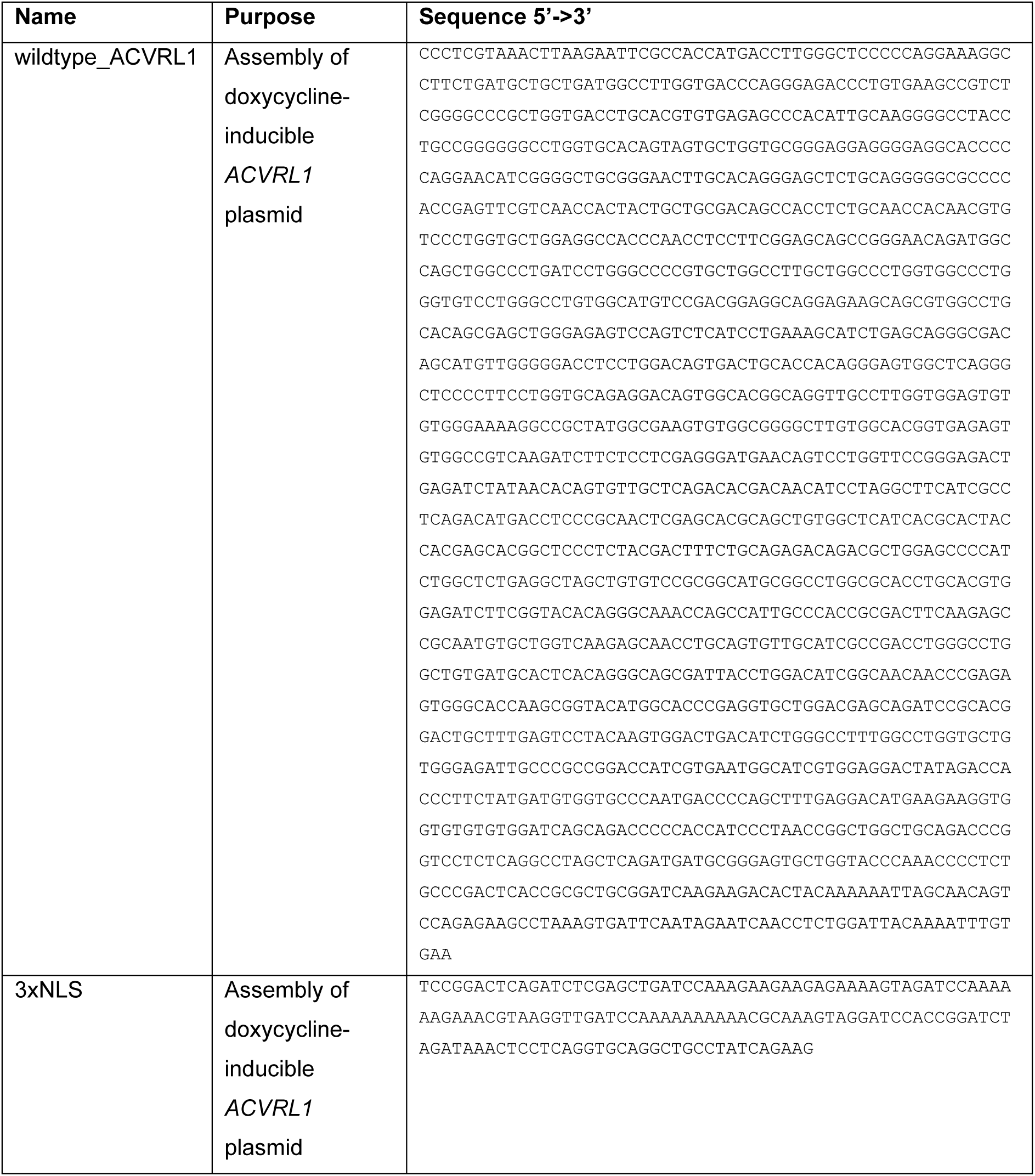
GeneBlocks.

**Table S3:** List of antibodies for flow cytometry (FC), IF, and IB.

| Antibody | Species | Dilution | Purpose | #RRID | Catalog # |
| --- | --- | --- | --- | --- | --- |
| NANOG_PE | Mouse | 1:5 | FC | AB_1645522 | 560483 |
| OCT3/4_BV421 | Mouse | 1:25 | FC | AB_2739320 | 565644 |
| SSEA-4_FITC | Human | 1:25 | FC | AB_2653517 | 130-098-371 |
| ALK1 | Goat | 1:50 | FC | AB_2222591 | #AF370 |
| pSMAD1/5 | Rabbit | 1:1000 | WB | AB_491015 | #9516 |
|  |  | 1:250 | IF |  |  |
| SMAD1 | Rabbit | 1:1000 | WB | AB_2107780 | #9743 |
|  |  | 1:250 | IF |  |  |
| VIN | Mouse | 1:1000 | WB | AB_1131294 | sc-73614 |
| CD31 | Sheep | 1:800 | IF | AB_355617 | AF806 |
| Alexa Fluor 488; anti-rabbit | Donkey | 1:400 | IF | AB_2535792 | A21206 |
| Alexa Fluor 488; anti-goat | Donkey | 1:1000 | FC | AB_2534102 | A11055 |
| Alexa Fluor 647; anti-sheep | Donkey | 1:400 | IF | AB_10374882 | A21448 |
| HRP-anti-goat | Donkey | 1:2000 | WB | AB_2534673 | A15999 |
| HRP-anti-mouse | Horse | 1:2000 | WB | AB_330924 | #7076S |
| HRP-anti-rabbit | Goat | 1:2000 | WB | AB_2099233 | #7074S |

**Table S4:**
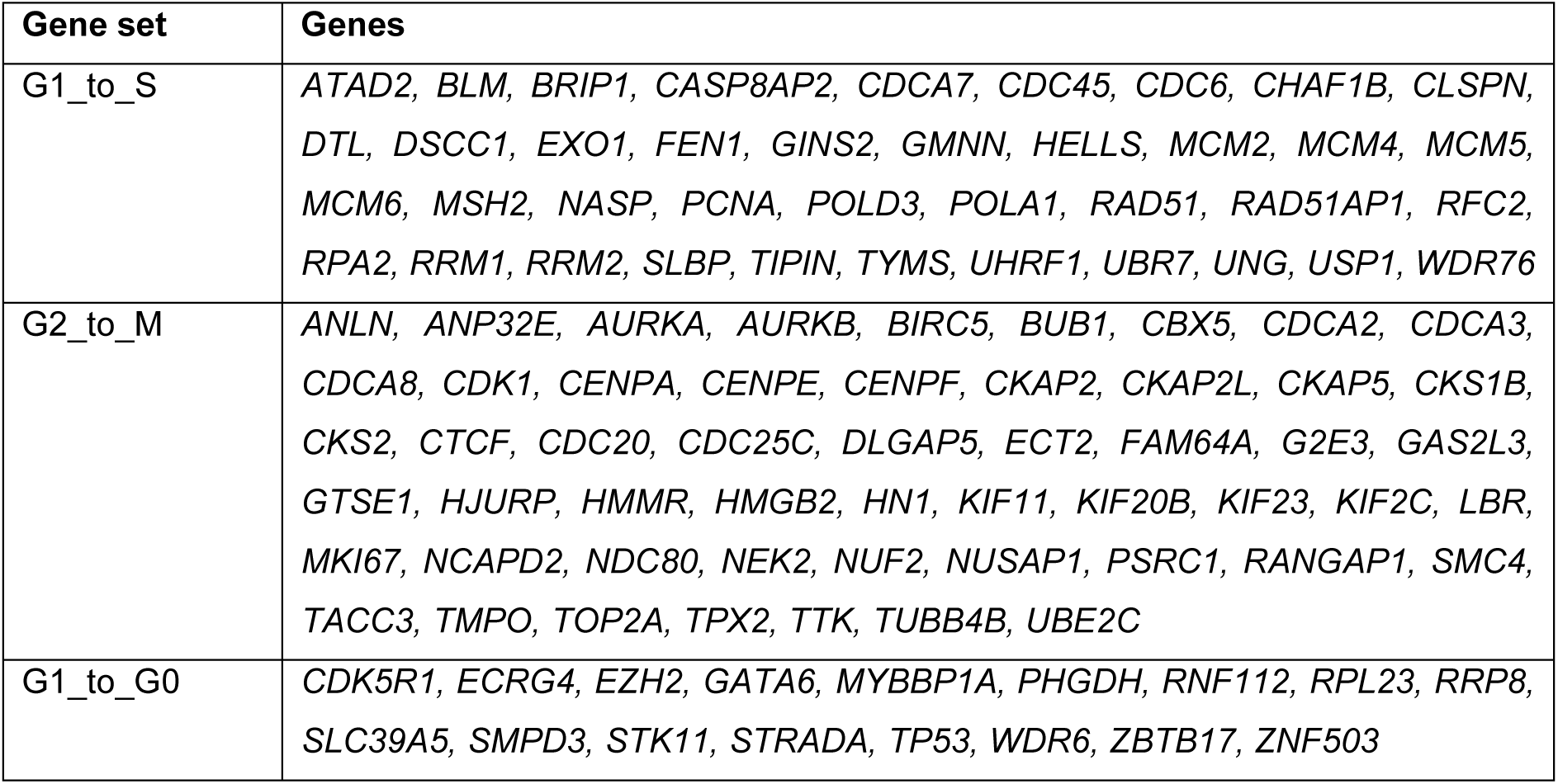
G1/S, G2/M, and G1/G0 gene sets.

| Gene set | Genes |
| --- | --- |
| G1_to_S | *ATAD2, BLM, BRIP1, CASP8AP2, CDCA7, CDC45, CDC6, CHAF1B, CLSPN, DTL, DSCC1, EXO1, FEN1, GINS2, GMNN, HELLS, MCM2, MCM4, MCM5, MCM6, MSH2, NASP, PCNA, POLD3, POLA1, RAD51, RAD51AP1, RFC2, RPA2, RRM1, RRM2, SLBP, TIPIN, TYMS, UHRF1, UBR7, UNG, USP1, WDR76* |
| G2_to_M | *ANLN, ANP32E, AURKA, AURKB, BIRC5, BUB1, CBX5, CDCA2, CDCA3, CDCA8, CDK1, CENPA, CENPE, CENPF, CKAP2, CKAP2L, CKAP5, CKS1B, CKS2, CTCF, CDC20, CDC25C, DLGAP5, ECT2, FAM64A, G2E3, GAS2L3, GTSE1, HJURP, HMMR, HMGB2, HN1, KIF11, KIF20B, KIF23, KIF2C, LBR, MKI67, NCAPD2, NDC80, NEK2, NUF2, NUSAP1, PSRC1, RANGAP1, SMC4, TACC3, TMPO, TOP2A, TPX2, TTK, TUBB4B, UBE2C* |
| G1_to_G0 | *CDK5R1, ECRG4, EZH2, GATA6, MYBBP1A, PHGDH, RNF112, RPL23, RRP8, SLC39A5, SMPD3, STK11, STRADA, TP53, WDR6, ZBTB17, ZNF503* |

